# Dendrimeric DNA Coordinate Barcoding Design for Spatial RNA Sequencing

**DOI:** 10.1101/2023.06.26.546618

**Authors:** Jiao Cao, Zhong Zheng, Di Sun, Xin Chen, Rui Cheng, Tianpeng Lv, Yu An, Junhua Zheng, Jia Song, Lingling Wu, Chaoyong Yang

**Affiliations:** Institute of Molecular Medicine, Shanghai Key Laboratory for Nucleic Acid Chemistry and Nanomedicine, Renji Hospital, School of Medicine, Shanghai Jiao Tong University, Shanghai 200127, China; The MOE Key Laboratory of Spectrochemical Analysis & Instrumentation, Department of Chemical Biology, College of Chemistry and Chemical Engineering, Xiamen University, Xiamen 361005, China; Department of Urology, Renji Hospital, Shanghai Jiao Tong University School of Medicine, Shanghai 200127, China

## Abstract

Spatially resolved transcriptomic technologies show promise in revealing complex pathophysiological processes, but developing sensitive, high-resolution, and cost-effective methodology is challenging. Here, we report a <u>de</u>ndrimeric DNA <u>co</u>ordinate barcoding <u>de</u>sign for spatial <u>R</u>NA <u>seq</u>uencing (Decoder-seq). This technology combined dendrimeric nano-substrates with microfluidic coordinate barcoding to generate high-density spatial DNA arrays with deterministically combinatorial barcodes in a resolution-flexible and cost-effective manner (∼$0.5/mm^2^). Decoder-seq achieved high RNA capture efficiency, ∼68.9% that of *in situ* sequencing, and enhanced the detection of lowly expressed genes by ∼five-fold compared to 10× Visium. Decoder-seq visualized a spatial single-cell atlas of mouse hippocampus at near-cellular resolution (15 μm) and revealed dendrite-enriched mRNAs. Application to renal cancers dissected the heterogeneous tumor microenvironment of two subtypes, and identified spatial gradient expressed genes with the potential in predicting tumor prognosis and progression. Decoder-seq is compatible with sensitivity, resolution, and cost, making spatial transcriptomic analysis accessible to wider biomedical applications and researchers.

---

Deciphering the spatial organization and molecular features of cells is crucial for comprehending various pathophysiological processes such as embryogenesis, neuroscience, and disease progression^1, 2^. Spatial transcriptomics technologies are powerful tools for delineating spatial gene expression patterns within tissues, revealing the composition of cell types, their spatial arrangements, and interactions^3, 4^. Existing technologies include two major methodologies: imaging-based *in-situ* hybridization (ISH) or *in-situ* sequencing (ISS), and spatially barcoded array-based sequencing^2^. Although the former provides subcellular resolution, it involves lengthy, repetitive, and technically-demanding imaging workflow^5–7^. Alternatively, spatially barcoded array-based methodologies encode tissue mRNAs at a large-scale using DNA arrays, enabling readout of mRNA type, abundance, and spatial locations through next-generation sequencing (NGS)^8^. They offer a high-throughput, straightforward approach to the whole-transcriptome analysis that has gained significant attention.

Several remarkable approaches have been developed to generate spatially barcoded DNA arrays for systematic spatially resolved transcriptomic profiling of tissue sections. They can be classified into deterministic and random barcoding strategies. The former approach typically leverages a micro-spotting method to deposit deterministic oligonucleotides (oligos) at specific positions on solid substrates^9^. However, the instrument’s precision limits the spot size to multicellular resolution (e.g., 55 μm diameter for each spot in commercial 10× Visium), and barcoding costs increase linearly with the number of spots. The latter uses randomly-generated DNA micro/nanoballs or clusters on substrates, yielding spatially barcoded DNA arrays with smaller features from cellular to subcellular resolution. This approach is used in technologies such as HDST^10^, Slide-seq^11^/Slide-seqV2^12^, Seq-scope^13^, Stereo-seq^14^, and Pixel-seq^15^. Random barcoding strategies require time-consuming and costly *in situ* sequencing to decode the DNA balls or clusters for position indexing. For example, in Seq-Scope, decoding the feature barcodes added $30/mm^2^ and 3-4 h per run^13^. In Pixel-seq, the cost of the crosslinked polyacrylamide “stamp gel” can be as low as $0.06/mm^2^, but the “sequenced gel” still costs $2.2/mm^2^ and 35 hours of sequencing time^15^. Thus, the currently available methods face challenges in balancing spatial resolution and cost.

Gene detection sensitivity is another challenge for spatially barcoded array-based approaches^16^, with ST having only 6.9% of the sensitivity of ISH^9^ and Slide-seq at 2.7%^11^. This may result in the omission of important low-abundance transcripts, restricting the range of biological problems that can be addressed with such methods. For example, charting spatial expression patterns of olfactory receptors (*Olfr*) is critical to understanding olfaction, but the lowly-expressed *Olfr* transcripts are difficult to sensitively detect by existing spatially barcoded array-based methods^17, 18^. Besides, current methods have limited capacity to detect molecular patterns in RNA-degraded precious clinical samples^19^, due to the reduced abundance of transcripts. In addition, the low detection sensitivity requires deeper sequencing depth to ensure enough gene coverage for analysis, increasing sequencing costs^20^. Obviously, the mRNA capture efficiency of DNA barcoded arrays determines the gene detection sensitivity of methods to a great extent. However, existing substrates suffer from low DNA modification density and molecular crowding during capturing, resulting in low mRNA capture efficiency.

Here, we report a <u>de</u>ndrimeric DNA <u>co</u>ordinate barcoding <u>de</u>sign for spatial <u>R</u>NA <u>seq</u>uencing (Decoder-seq), which is compatible with detection sensitivity, spatial resolution, and cost. This technology combines dendrimeric nano-substrates with microfluidic coordinate barcoding to generate deterministic *X_i_Y_j_*coordinate arrays of high-density DNA in a resolution-flexible and cost-effective manner. First, three-dimensional (3D) nano-substrates afford density enhancement of modified spatially DNA barcodes by ∼ an order of magnitude compared with reported spatial transcriptomics methods. The resultant high RNA capture efficiency of Decoder-seq, ∼68.9% compared to ISS, allows detection of lowly expressed *Olfr* genes with ∼5-fold higher than 10× Visium. Second, inspired by the idea of using an X-Y coordinate system to effectively define any location within a plane and also by the elegant DBiT-seq design of Fan’s group^21^, we adopted a microfluidics-assisted coordinate barcoding strategy to generate *X_i_Y_j_* coordinate DNA arrays with exceptional flexibility and precision in spatial resolution (50, 25, and 15 μm). Third, combinatorial barcoding of deterministic DNA *X_i_* and *Y_j_* reduces the variety of DNA barcodes and avoids decoding steps, dramatically decreasing array fabrication cost to ∼$0.5/mm^2^, compared favorably to similar methodologies developed to date. Decoder-seq of 15 μm resolution achieves transcript counts of 40.10 unique molecular identifiers (UMIs) detected per μm^2^ on average, outstanding among available spatial transcriptomics methodologies. It can dissect the fine structure of the mouse brain at a near-cellular resolution, thereby identifying low-expressed dendritic-enriched genes in neurons of the hippocampus. Application to renal cell carcinomas (RCC) dissects spatial architecture heterogeneity and commonalities between clear cell renal cell carcinomas (ccRCC) and chromophobe RCC (chRCC) subtypes for the first time. Epithelial-mesenchymal transition (EMT)-related genes with a spatial gradient pattern were identified as an important predictor of clinical stages and outcomes in RCCs.

## Results

### Overview of Decoder-seq

Decoder-seq mainly includes three parts: the generation of 3D dendrimeric DNA coordinate barcoding array, spatial indexing of tissue transcriptome, and spatially resolved transcriptomics analysis (**Figure 1**). We first assembled 3D dendrimer monolayers on glass slides. Each spherically-shaped dendrimer had 64 active primary amino groups, enabling covalent cross-linking of high-density spatial DNA barcodes and facilitating efficient mRNA capture. A pair of microchannel chips perpendicular to each other were then designed and successively placed on the 3D dendrimeric slide to introduce sets of barcode solutions. Through the covalent linkage of DNA barcodes *X_1_*-*X_n_*with dendrimeric substrates, and subsequent ligation to DNA barcodes *Y_1_*-*Y_m_*, deterministic barcoded DNA arrays were formed at the intersections with combinatorial *X_i_Y_j_* coordinates from barcodes *X_i_* and *Y_j_*. Here, DNA barcode X consists of a 5’ end amino modification, a PCR handle, a distinct 8-base barcode X index, and a linker 1 for ligation reaction to introduce the second barcode Y. DNA barcode Y consists of a linker 2, a unique 8-base barcode Y index, a UMI, and an oligo-dT sequence for mRNA capture (**Figure S1**). With this combinatorial barcoding, a spatially barcoded array with n × m spots required only n + m types of DNA barcodes, rather than the n × m unique DNA barcodes required in traditional barcoding strategies (**Figure 1a**). Next, a tissue section was mounted onto the barcoding array for spatial transcriptome indexing. Following tissue fixation, hematoxylin and eosin (H&E) staining, and permeabilization, mRNAs were captured by oligo-dT sequences on slides for *in situ* reverse transcription (RT) (**Figure 1b**). Finally, after amplification, library preparation, and paired-end sequencing, the sequences containing both cDNA and *X_i_Y_j_* coordinates were analyzed for spatially resolved transcriptome mapping (**Figure 1c**). Decoder-seq, a microfluidics-assisted dendrimeric DNA coordinate barcoding design, is therefore expected to be compatible with high sensitivity, high spatial resolution, and low cost.

**Figure. 1.**
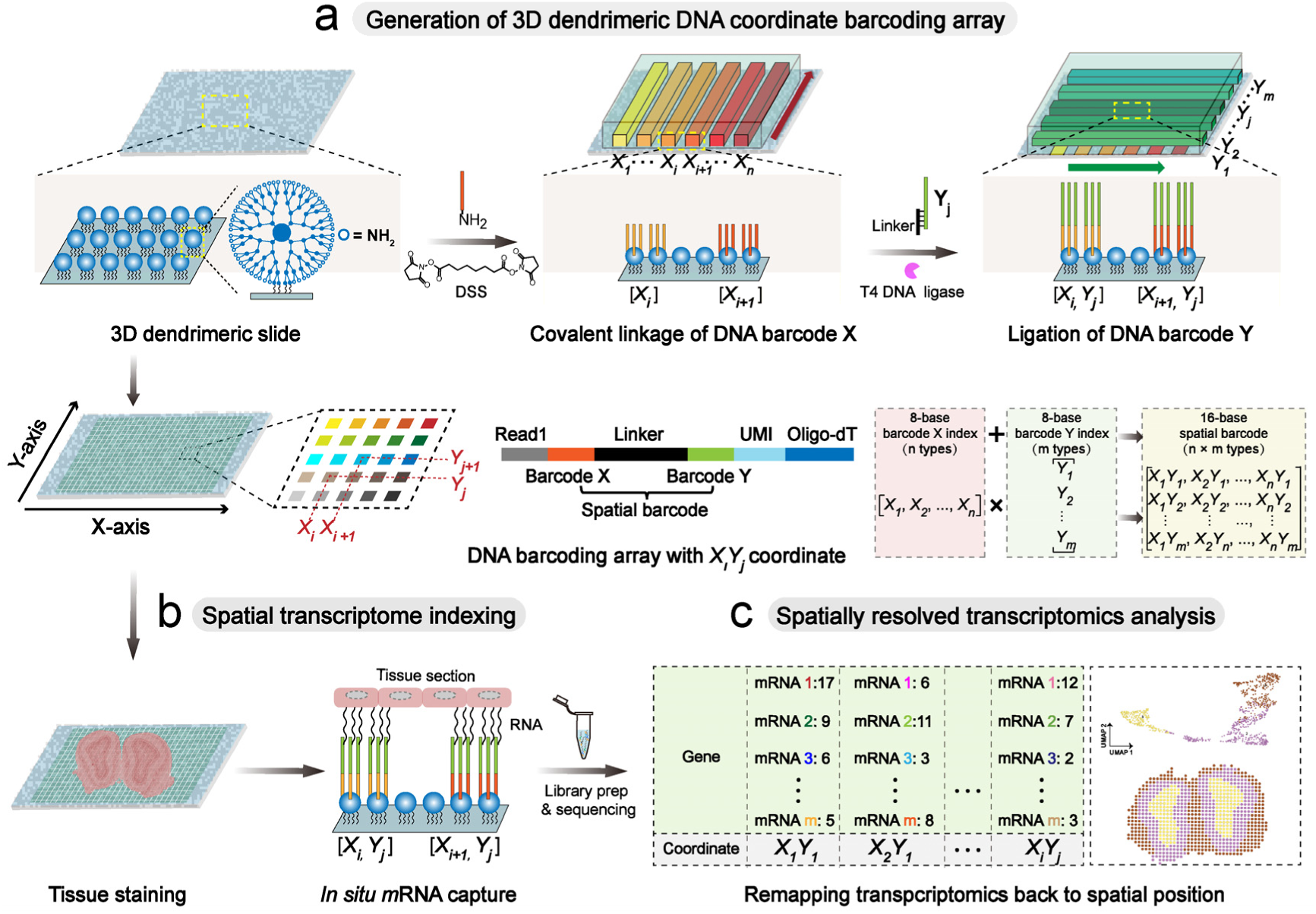
Decoder-seq workflow. (**a**) Generation of 3D dendrimeric DNA coordinate barcoding array. 3D dendrimeric nano-substrates offer multiple active primary amino groups to allow high-density DNA modification. A pair of microchannel chips consecutively introduce two sets of DNA barcodes to dendrimeric nano-substrates, allowing covalent linkage of DNA barcode X and ligation of DNA barcode Y to form *X_i_Y_j_* coordinates at the intersections. The linker consists of linker 1 and linker 2. Linker 1 from 3’ DNA barcode X while linker 2 from 5’ DNA barcode Y. Thus, a DNA barcoding array with n × m unique *X_i_Y_j_* coordinates was formed with only n + m unique DNA barcodes. **(b)** Spatial transcriptome indexing. *In situ* capture and RT of mRNAs from tissue sections were conducted on DNA coordinate barcoding arrays, resulting in cDNAs for library preparation and sequencing. **(c)** Spatially resolved transcriptomics analysis. A spatially-resolved transcriptomics map was reconstructed by integrating gene expression data with the spatial barcodes via bioinformatic analysis.

## 3D dendrimeric nano-substrates enable modification of high-density DNA barcodes

For highly sensitive and accurate spatial transcriptomics sequencing, it is desirable to modify spatial DNA barcodes onto a substrate with high density, uniform distribution, and increased accessibility for efficient mRNA capture. To this end, we fabricated 3D dendrimeric substrates for spatially barcoded arrays generation by assembling poly(amidoamine) (PAMAM) G4 dendrimers on glass slides (**Figure 1a, Figure S1**). A two-dimensional (2D) planar epoxy-modified slide was fabricated through a silanization reaction, which was then reacted with dendrimers via a ring-opening process to create a 3D dendrimeric slide (**Figure S1a**). With disuccinimidyl suberate (DSS), a homobifunctional reagent with double-terminal N-HydroxySuccinimide (NHS) groups, 5’ end amino of DNA barcode X was cross-linked onto dendrimeric substrates rapidly and conveniently (**Figure S1b**). Through T4 DNA ligase and linker sequences, DNA barcode X was ligated to DNA barcode Y to form the combinatorial spatial barcoding of *X_i_Y_j_* (**Figure S1c**).

Fluorescence images showed obvious fluorescence signals on the 3D dendrimeric slide upon reaction with amino-functionalized fluorescent DNAs (**Figure S2a**). In contrast, fluorescence signals were barely detected from the slide for fluorescent DNAs lacking primary amino modifications, indicating that the inert amino groups on cytosine and adenine bases did not undergo covalent linkage (**Figure S2b**). These results verified that the 3D dendrimeric slide had successfully modified DNA barcode X. Similarly, fluorescence imaging confirmed the successful ligation of DNA barcode Y (**Figure S2c-d**).

The modification of spherical dendrimers provides nano-substrates over the planar glass surface, increasing available surfaces for DNA modification. As demonstrated with atomic force microscopic (AFM) imaging, the assembly of PAMAM G4 (which has a diameter of ∼4.5 nm^22^) led to a significant increase in surface roughness (**Figure 2a-i, ii**). This result indicated that the dendrimeric surface offered nanostructured substrates. In addition, each dendrimer possessed 64 active primary amino groups on the periphery, facilitating the high-density modification of spatial DNA barcodes. As expected, DNA modification significantly increased surface roughness (**Fig. 2a-iii**), while no obvious change was observed on the 2D slides (**Figure S3**), indicating a higher density of modified DNA barcodes on the 3D dendrimeric slides. Fluorescence imaging analysis verified much higher densities of modified DNA barcodes on 3D slides than those on 2D slides (**Figure S4**). There were calculated to be ∼2.04 × 10^5^ modified spatial DNA barcodes per μm^2^ on the 3D dendrimeric slides, which is ∼ one order of magnitude higher than the number of modified barcodes on the 2D planar slides and other substrates used in reported spatial transcriptomics methods^9, 15^ (**Figure 2b, Table S1, Methods**). Although the spatial DNA barcodes were generated by a two-step reaction in Decoder-seq, the high modification density could be achieved with the mediator of the 3D nanostructured dendrimers on slides. To validate the ligation-based combinatorial spatial barcoding strategy, TA clone sequencing was utilized to identify spatial barcodes (8-base barcode X + 8-base barcode Y). As high as 92.7 ± 1.2% of the modified oligos exhibited the expected base sequence, demonstrating the feasibility and reliability of this strategy for generating coordinate DNA arrays (**Figure 2c**).

**Figure. 2.**
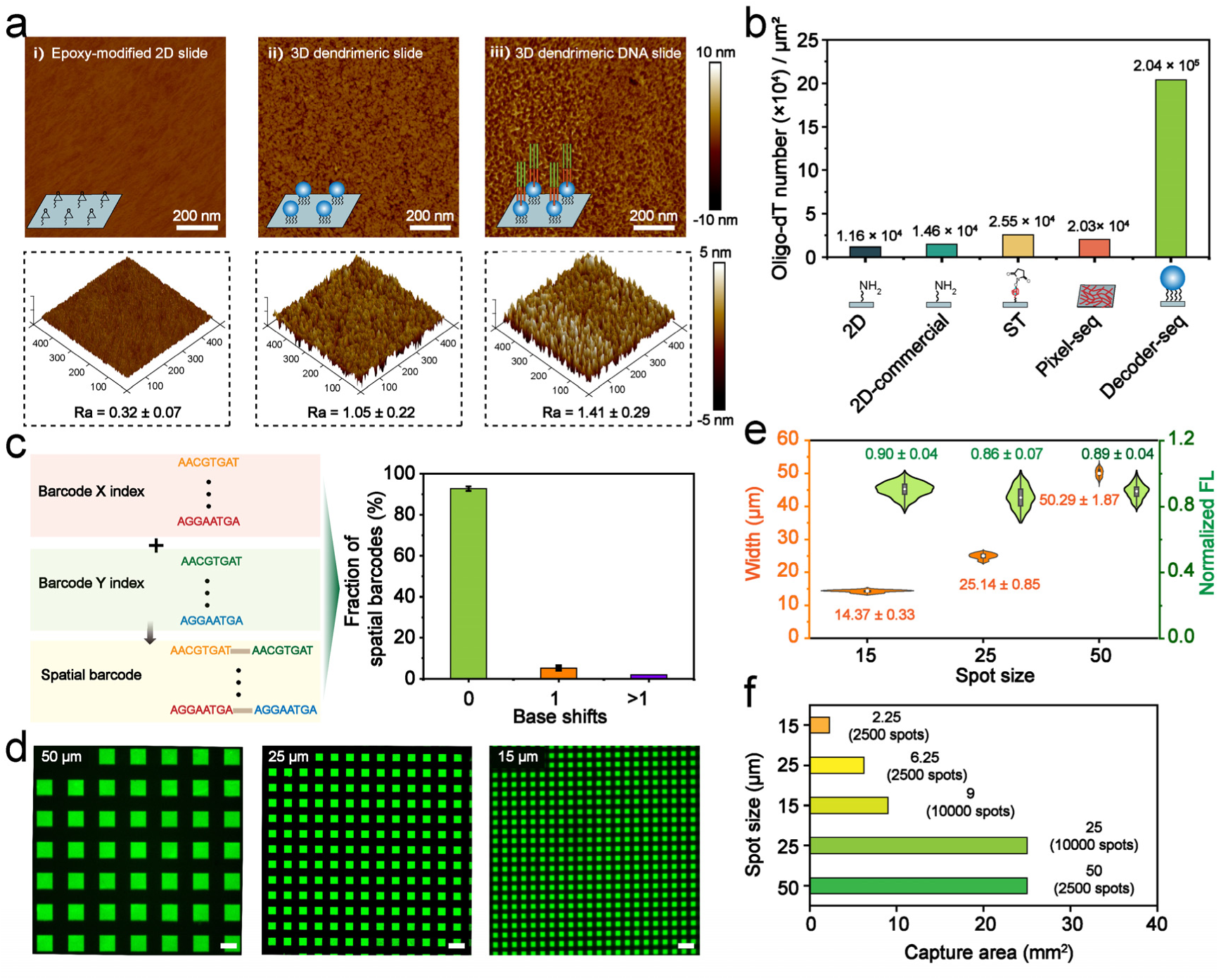
Characterization of 3D dendrimeric DNA coordinate barcoding arrays. (**a**) AFM imaging to characterize the surface morphology of the following functionalized slides: i) epoxy-modified 2D slide, ii) 3D dendrimeric slide, and iii) 3D dendrimeric DNA slide. Top: AFM height images; Bottom: 3D AFM images of the corresponding slides. Ra: average roughness. **(b)** Comparison of the spatial DNA barcodes (oligo-dT) density on 2D amino slides generated for this study (**Methods**), 2D commercial amino slides, 3D dendrimeric slides (Decoder-seq), and other substrates used in ST^9^ and Pixel-seq^15^. **(c)** The accuracy of spatial barcodes was verified with TA clone sequencing (n = 3). 92.7 ± 1.2% of spatial barcodes showed no base shifts, deletions, additions, or substitutions. **(d)** Fluorescence images of 3D dendrimeric DNA coordinate barcoding arrays consisting of 50, 25, and 15 μm spots. Alexa 488-labeled DNA barcode Y was ligated to the immobilized DNA barcode X at the intersections of the first set (*X_1_*–*X_50_*) and the second set (*Y_1_*–*Y_50_*) of flows, generating square fluorescent spots. Scale bar: 50 μm. **(e)** Spot size and fluorescence intensity quantification. The measured spot sizes were consistent with the microchannel sizes. The fluorescence intensities indicated uniform DNA modification. **(f)** The array capture areas corresponding to different spot numbers and sizes.

### Microfluidics-assisted spatial barcoding of dendrimeric DNA coordinate arrays

The high cost of current spatial transcriptomics methods precludes their wider application in scientific and clinical research. The cost is mainly governed by the fabrication of spatially barcoded arrays, such as requiring thousands of unique DNA barcodes or costly spatial decoding processes. To reduce costs and simplify the fabrication process, we developed a microfluidics-assisted coordinate barcoding strategy (**Figure 1a, Figure S5a-b**). Using customized clamps, a pair of microchannel chips could be sequentially held on the dendrimeric slide firmly without liquid leakage (**Figure S5c-d**). This enabled the delivery of deterministic DNA barcode X and Y reagents onto a slide to generate spatially barcoded DNA arrays with coordinates of combinatorial DNA *X_i_Y_j_* at the intersections.

Based on the combinatorial barcoding strategy, the number of unique DNA barcodes required is greatly reduced from n × m to n + m. For example, 10,000 (100 × 100) spatially barcoded spots can be generated using only 200 spatial DNA barcodes. The hyperbranched and flexible 3D dendrimeric substrates, and the covalent linkage strategy, dramatically reduced both the DNA reaction concentration (as low as 5 μM) and reaction time (∼40 min) compared with the DNA modification strategy used in ST^23^ (33 μM, incubation overnight). Thus, our approach reduced the required variety and concentration of DNA and generated deterministic *X_i_Y_j_* coordinates that were decoding-free. Combined, these factors contributed to the much lower cost of the dendrimeric DNA coordinate barcoding array compared with other spatially barcoded arrays. For example, ignoring differences in instrumentation requirements and their associated costs, each array cost only $0.5/mm^2^ (**Table S2-3**), 70 times less than the DNB chips used in Stereo-seq^14^ ($35/mm^2^), and 300 times cheaper than the Illumina flow cells used in Seq-scope^13^ ($150/mm^2^). The cost could be further reduced with multiplexing encoding on slides^24^.

Tissue types vary in size, and typically comprise various cells with diameters of ∼10 μm. It is highly desirable to fabricate spatially barcoded arrays with flexibly barcoded areas and high spatial resolution. To achieve this, we designed microchannel chips that had different channel widths and numbers, providing exceptional flexibility and precision in the spatial resolution (spot size) and capture area for spatially barcoded arrays (**Figure S5a**). As a proof of concept, three kinds of DNA coordinate arrays with spot widths of 50, 25, and 15 μm were generated on 3D dendrimeric slides (**Figure 2d**). These exhibited satisfactory uniformity of spot size and fluorescence intensity, guaranteeing their accuracy and impartiality for subsequent spatial transcriptomics sequencing (**Figure 2e**). Moreover, the 15 μm diameter spot size would theoretically enable near-single-cell spatial transcriptomics analysis. We also fabricated spatially barcoded arrays with varying capture areas and spot widths to match different tissue sizes (**Figure 2f, Figure S6**), such as the mouse hippocampus (∼2.5 mm^2^), mouse olfactory bulb (MOB, ∼10.5 mm^2^), and mouse hemibrain (∼24.2 mm^2^).

### Dendrimeric DNA coordinate barcoding arrays capture spatial mRNAs of tissues with high efficiency

We evaluated the RNA-capturing capability of the dendrimeric DNA substrate using a Cy3-dCTP-mediated cDNA labeling assay (**Methods**). MOBs were selected as model tissues due to their distinct layered structure and well-defined molecular signatures. The dendrimeric DNA substrate generated a clear cDNA fluorescence pattern that well matched the tissue structure revealed by general histology (**Figure 3a**). Fluorescent cDNAs were strictly localized under tissue cells compared with tissue DNA staining (**Figure S7**). The mRNA lateral diffusion distance was measured at 1.6 ± 0.7 μm, similar to ST (1.7 ± 2 μm)^9^. In addition, we used the signal intensity of fluorescent cDNA patterns as an index to visually evaluate tissue mRNA capture performance on different substrates. Fluorescent cDNA signals were significantly stronger on the 3D dendrimeric DNA substrates than on the 2D planar DNA substrates (**Figure S8**), indicating a higher mRNA capture efficiency.

**Figure. 3.**
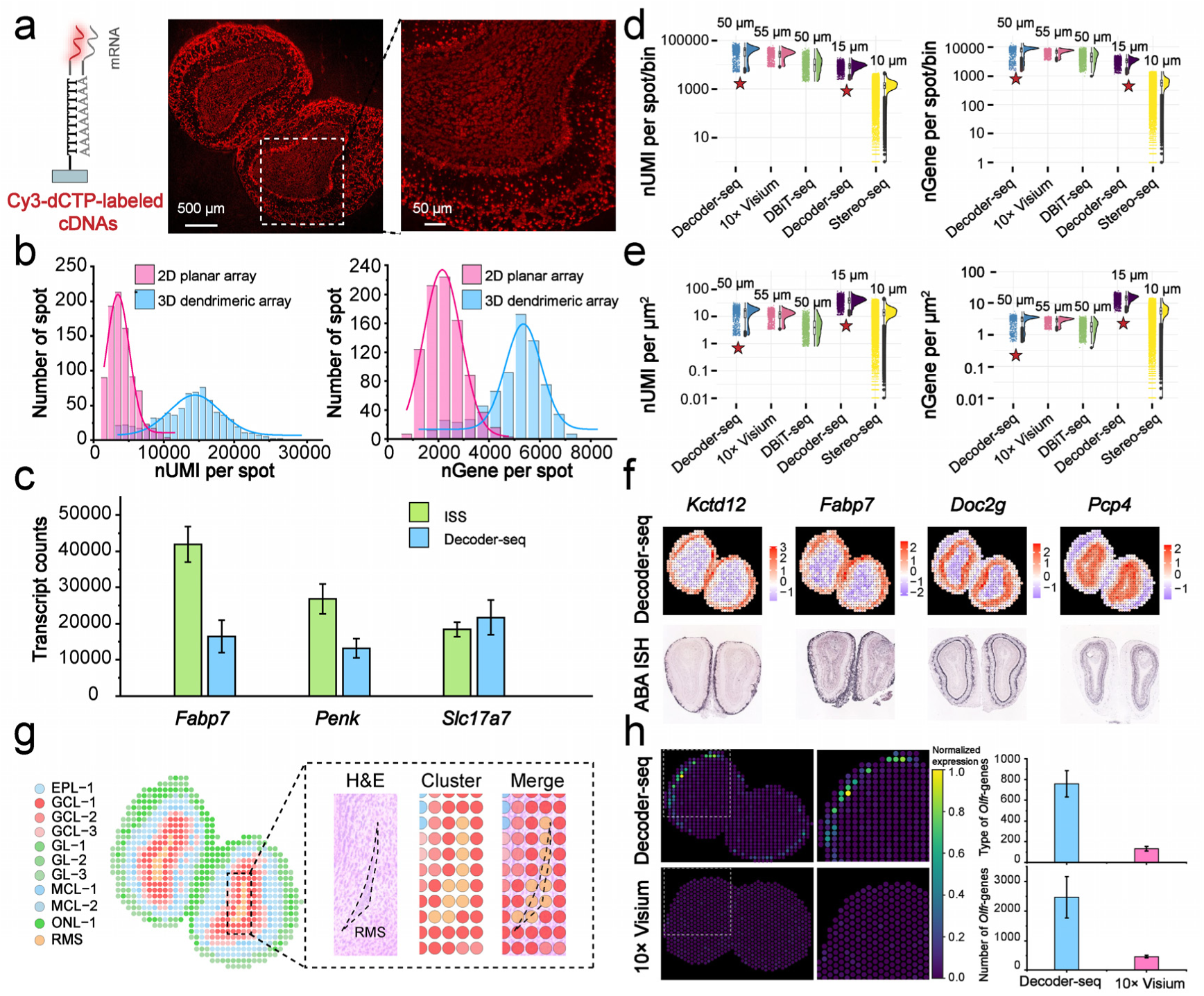
Spatially resolved transcriptomes analysis of MOBs with Decoder-seq. (**a**) Fluorescent cDNA footprints of the MOB on a 3D dendrimeric slide, in which cDNA was labeled by Cy3-dCTP. Left: representation of a labeled cDNA molecule. Middle: fluorescent cDNA image. Right: magnified view of the region outlined in white in the middle image. **(b)** Comparison of UMI (left) and genes (right) detected in two MOB sections with a 2D planar DNA coordinate barcoding array and 3D dendrimeric DNA coordinate barcoding array, respectively. nUMI: number of UMIs; nGene: number of genes. **(c)** Comparison of the number of transcripts detected with Decoder-seq and ISS corresponding to three selected genes. Transcript counts were calculated from adjacent MOB sections (n = 3). **(d)** Distribution of gene and UMI counts per spot/bin. Decoder-seq was compared to 10× Visium, DBiT-seq, and Stereo-seq with different spot/bin sizes. Fresh frozen MOB tissues were used in Decoder-seq, 10× Visium, and Stereo-seq^14^. Formaldehyde-fixed mouse embryo tissue was used in DBiT-seq^21^. **(e)** Normalized nUMI and nGene per μm^2^ for each method. **(f)** Spatial visualization of *Kctd12*, *Fabp7*, *Doc2g*, and *Pcp4* expression in MOB sections and the corresponding ISH images of the adult MOB taken from ABA^25^. **(g)** Unsupervised clustering of the MOB section detected by Decoder-seq at 50 μm spatial resolution (left), and magnified views of RMS regions (right, black dotted box). ONL, olfactory neuron layer; GL, glomerular layer; EPL, external plexiform layer; MCL, mitral and tufted cell layers; GCL, granule cell layer; RMS, rostral migratory stream. **(h)** Spatial expression (left) and comparison of the types (upper right) and numbers (lower right) of *Olfr* genes detected with Decoder-seq and 10× Visium.

Encouraged by this, we profiled MOBs to assess the performance of Decoder-seq. We assayed 10 μm MOB sections using dendrimeric DNA coordinate barcoding arrays with 50 μm and 15 μm resolution. Sequencing libraries were constructed for paired-end sequencing to identify spatial barcodes and the mRNA expressions at each spot (**Figure S9 and S10a-b**). In the 50 μm-spot Decoder-seq experiments, an average of 35,720 UMIs and 7,436 genes were detected per spot with a sequencing depth of 76% on the MOB library. The gene and UMI distributions between the two replicates were highly consistent (R^2^ = 0.96) (**Figure S10c-d**). At the same sequencing depth (40M reads), Decoder-seq outperformed the dendrimer-free 2D DNA coordinate barcoding array, detecting more UMIs and genes per spot (**Figure 3b**). This demonstrated the high-density spatial DNA barcodes afforded by Decoder-seq facilitated efficient mRNA capture, resulting in high-sensitivity gene detection.

Next, Decoder-seq was validated with ISS to assess its mRNA capture efficiency. Specific genes (*Fabp7*, *Slc17a7*, and *Penk*) in Decoder-seq were quantified and compared to the corresponding counts detected by ISS from adjacent MOB sections (**Figure 3c, Table S4**). Decoder-seq was calculated to detect an average of 68.9 ± 15.6% of total mRNA transcripts defined by ISS. Previous research has quantified the detection sensitivity of ISS to be ∼30% of available transcripts in cells^26^. Thus, we estimated the overall detection sensitivity of Decoder-seq to be ∼20.68%, which surpassed existing spatial transcriptomics methods, such as ST^9^ (∼6.9%), and DBiT-seq^21^ (∼15.5%).

The capability of Decoder-seq for detecting UMIs and genes was further evaluated by comparison with existing spatial transcriptomics methods (**Figure 3d-e**). For 50 μm-spot Decoder-seq, the per-spot UMIs and genes count outperformed those of 50 μm-pixel DBiT-seq and 55 μm-spot 10× Visium. Notably, Decoder-seq detected significantly more genes than 10× Visium, with an increase of ∼ 122% based on saturation curves (**Figure S10e-f**). More importantly, Decoder-seq could possess both high spatial resolution and high gene detection sensitivity. With improved spatial resolution, 15 μm-spot Decoder-seq detected an average of 9,024 UMIs and 3,302 genes per spot (**Figure 3e**). Excellent reproducibility was also observed between replicates (**Figure S10g**). Considering the different spot/bin sizes between these technologies, we normalized the transcriptome capture performance to the array unit area (μm^2^). Decoder-seq at both 50 μm and 15 μm resolution was superior to other technologies, showing a higher number of UMIs and genes detected per μm^2^ (**Figure 3e**). Remarkably, 15 μm-spot Decoder-seq achieved counts of 40.1 UMIs and 14.7 genes per μm^2^, which was significantly higher than other methods with cellular/subcellular resolution such as Stereo-seq^14^ (UMI:14.5, gene: 5), pixel-seq^15^ (UMI:9.8), Seq-scope^13^ (UMI:10), slide-seqV2^12^ (UMI: 6.2) and HDST^10^ (UMI: 0.12). Collectively, Decoder-seq provides an outstanding mRNA capture efficiency and thus sensitive gene detection.

### Decoder-seq achieves accurate spatial gene-expression patterns and high-sensitive detection of lowly-expressed genes

Decoder-seq demonstrated the accurate spatial distribution of specific genes (*Kctd12, Fabp7*, *Pcp4*, *Doc2g*) and exhibited expected similarity to that of ISH data taken from Allen Brain Atlas (ABA)^25^ (**Figure 3f**). This accuracy was further validated by ISS for six specific genes (*Kctd12*, *Doc2g*, *Pcp4*, *Fabp7*, *Slc17a7*, *and Penk*) (**Figure S11**). We observed clearer spatial patterns of gene expression at near-cellular (15 μm) resolution, such as spatial distribution of *cdhr1* and *Eomes* in the mitral and tufted cell layers (MCL) (**Figure S12**).

We performed unsupervised clustering of all spots using the mRNA expression profiles and identified eleven distinct clusters. We annotated clusters based on region-specific marker genes and spatial locations (**Figure S12a**), including the olfactory nerve layer (ONL), rostral migratory stream (RMS), granular layer (GCL), MCL, external plexiform (EPL) and glomerular layer (GL). The spatial distribution of these clusters recapitulated the layer-specific patterns, consistent with well-defined layered anatomical structures (**Figure 3g, left**). For example, the RMS cluster was resolved as a single layer of spots, consistent in size with the width observed in the morphological layer, indicating the excellent spatial resolution and minimal mRNA lateral diffusion of Decoder-seq (**Figure 3g, right**). Moreover, Decoder-seq allowed for tissue organization profiling at a higher spatial resolution. This was demonstrated by the spatial distribution of identified clusters using 15 μm-spot Decoder-seq, which revealed finer structures that were correlated with tissue morphology (**Figure S12b**). A combination of 50 μm-spot and 15 μm-spot Decoder-seq would enable large field-of-view spatial transcriptomics and further reveal near-cellular resolution in selected regions.

Olfactory sensory processing is organized across olfactory bulb glomeruli, where axons of peripheral sensory neurons (PSNs) expressing the same *Olfr* converge on invariant glomeruli to transmit odorants-specific activity to central neurons^17, 18^. The mammalian olfactory receptor repertoire consists of ∼ 1,000 *Olfr* genes, but ISH-based studies have indicated that each cell expresses only one or a few of this genes^27^. Revealing the distribution of *Olfr* genes within glomeruli is critical to understanding odor perception, whereas high-throughput analyses have been hindered by the lowly-expressed *Olfr* genes. For example, ST only detected a mere 77 *Olfr* genes^9^. With the high detection sensitivity and exquisite spatial resolution of Decoder-seq, we were encouraged to examine the lowly-expressed *Olfr* genes in MOBs. 50 μm-spot Decoder-seq identified 2,470 UMIs and 731 genes of *Olfr*, remarkably higher than those detected by 10× Visium (**Figure 3h, Table S5-6**). Moreover, the spatial distribution of *Olfr* genes was observed in the expected layers of the GL. Taken together, Decoder-seq achieves accurate spatial gene-expression patterns and high-sensitive detection of lowly-expressed genes.

### Decoder-seq enables near-cellular spatial transcriptomics analysis

Dendritic-enriched mRNAs in hippocampal neurons are implicated in synaptic plasticity and learning^28, 29^. However, they constitute only a small fraction of neuronal mRNAs, most of which are located in somata^30^ (**Figure 4a**), and identification of their spatial distribution requires methods with high sensitivity and resolution. Due to the high sensitivity and near-cellular resolution of 15 μm-spot Decoder-seq, this method was further explored to map dendritic-enriched mRNAs in mouse hippocampus. Compared with the 50 μm-spot Decoder-seq data (**Figure S13a**), the total UMI count map derived from 15 μm-spot Decoder-seq exhibited finer structures that were correlated with hippocampal morphology (**Figure 4b**). A drop in the total UMIs numbers from CA1 somata to stratum radiatum occurred over a distance matching the expected size of somata (∼50 μm).

**Figure. 4.**
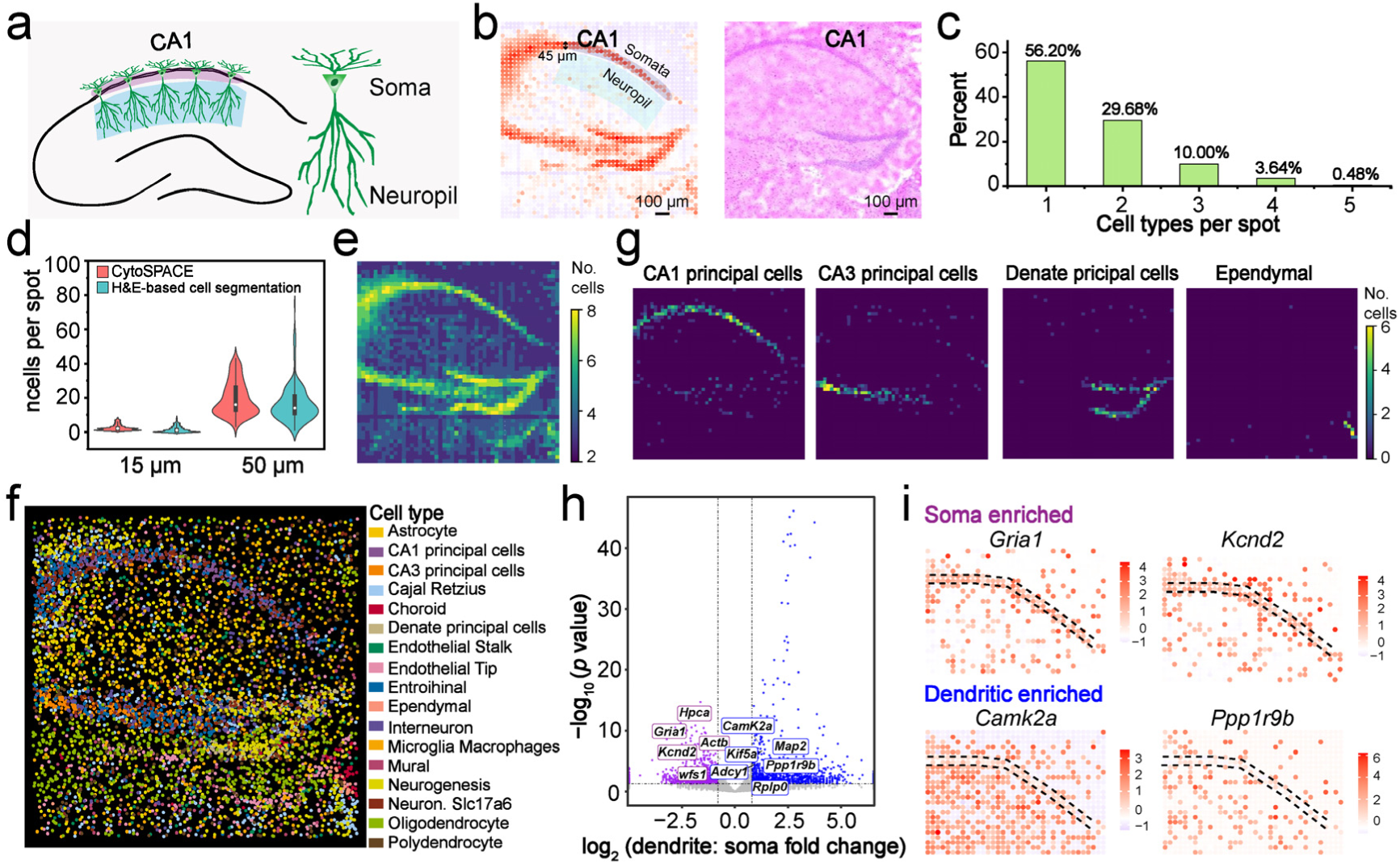
Decoder-seq resolves the mouse hippocampus at near cellular resolution. (**a**) Schematic diagram of neurons in the CA1 region of mouse hippocampus. **(b)** Spatial distribution of UMIs in the hippocampus (left) and the H&E image of the mouse hippocampus (right). **(c)** The number of cell types assigned per 15 μm-spot. **(d)** Quantification of the number of cells per spot. Cell numbers per spot were calculated using H&E-based cell segmentation and Cytospace. ncells: number of cells. **(e)** Spatial distribution of total cells in the hippocampus. **(f)** A spatial single-cell atlas of the mouse hippocampus. **(g)** Regional enrichment of CA1 principal cells, CA3 principal cells, dentate principal cells, and ependymal cells. **(h)** Differentially expressed genes between the somata versus proximal dendrites in the CA1 region. Two-tailed, two-sample t-tests were performed and genes with a false discovery rate (FDR)-corrected P value of <0.05 and a fold change >0.08 are highlighted. **(i)** Spatial expression of soma-enriched and dendrite-enriched genes.

To investigate whether 15 μm-spot Decoder-seq enabled near-cellular spatial transcriptomics analysis, we applied CytoSPACE^31^ to predict the number and types of cells in each spot by mapping cells from a single-cell RNA sequencing (scRNA-seq) atlas onto the Decoder-seq data. We observed that 56.20% of the 15 μm-spots matched a single cell type, and 29.68% contained two distinct cell types (**Figure 4c**). In contrast, only 25.4% of spots from the 50 μm array showed mRNA from a single cell type (**Figure S13b**). The image-based cell segmentation algorithm that identified nuclei on H&E images further revealed a near-cellular resolution of this method (**Figure S13c, Methods**). With both algorithms, it was calculated that each 15 μm-spot contained an average of 2.5 ± 1.8 cells and 1.5 ± 1.9 cells, respectively, while each 50-μm – spot contained 19.6 ± 9.9 and 16.9 ± 10.8 cells, respectively (**Figure 4d**). Heat maps of cell numbers in spatial distribution visually demonstrated finer spatial structures of the hippocampus using Decoder-seq of 15 μm-spot resolution compared with that of 50 μm-spot resolution (**Figure 4e and Figure S14-15**). Overall, Decoder-seq with 15 μm spots demonstrated near cellular resolution and potential for finer-scale spatial analysis.

We used CytoSPACE to generate a spatial single-cell atlas that recapitulated the spatial distributions of 17 classical neuronal and non-neuronal cell types, including CA1 principal cells, CA3 principal cells, dentate principal cells, microglia, astrocytes, etc. (**Figure 4f**). The regional enrichment of specific cell types and their identified marker genes corresponded to the anatomical structure of the hippocampus (**Figure 4g, Figure S16a**). For example, CA1/CA3 principal cells were predominantly abundant in the Cornu Ammonis (CA) subfield on the“cord-like” structure of the section. Notably, the fine spatial features of the single-cell ependymal cell layer were also resolved (**Figure 4g**). We then used Decoder-seq to identify dendritically localized mRNAs within CA1 neuronal cells. Based on the stereotypical architecture of CA1 neuropil, we simplified transcript localization to a one-dimensional profile perpendicular to the CA1 soma layer (from the stratum pyramidale to the stratum radiatum; **Figure 4b**). Using scRNA-seq data of the hippocampus, we excluded non-neuronal markers and identified ∼301 genes with greater than twofold dendritic enrichment using differential gene expression analysis between the proximal neuropil (stratum radiatum) and soma (stratum pyramidale) (**Figure 4h, Table S7**). The significant overlap (p < 10^−1^^6^, hypergeometric test; **Figure S16b, Table S7**) between these genes and previously identified dendritically localized mRNAs suggests that Decoder-seq has the potential to identify dendritic-enriched genes^12, 32–34^. The spatial distribution of several known soma-enriched genes and dendritic-enriched genes were exhibited in **Figure 4i**. For instance, the synaptic plasticity-associated gene *Camk2a* was the most abundant in dendrites, and the *Ppp1r9b* was present in distal regions of dendrites, consistent with previous studies^6, 35^. These data demonstrated the sensitivity and robustness of Decoder-seq for identifying dendritic-enriched genes.

### Decoder-seq resolves spatial heterogeneity in RCCs

Tumors exist in a spatially structured ecosystem, the tumor microenvironment. TMEs exhibit significant heterogeneity in cell composition and spatial distribution, which are closely related to tumor biology and coordinated with tumor development^36^. A spatial transcriptomics technology with high sensitivity would enable a comprehensive and precise analysis of this complex architecture. We therefore applied Decoder-seq to decipher the spatial architecture of the TME in two RCC subtypes: ccRCC and chRCC. To date, only a few investigations have compared the differences in TMEs between ccRCC and chRCC subtypes at the single-cell level^37–39^. We collected eight primary biopsy tissues from five RCC patients (three with ccRCC and two with chRCC) and conducted 50 μm-spot Decoder-seq experiments (**Figure 5a, Methods**). Two regions, the leading-edge (L) and tumor core (T), were selected for each sample except ccRCC-3 and chRCC-2, which only had the L section due to extensive necrosis within the tumor core. Decoder-seq demonstrated exceptional gene detection sensitivity, with an average of 3,319 genes per spot. Even in sample regions with RNA degradation below the requirements for spatial transcriptomics analysis (RNA integrity number (RIN) < 7), Decoder-seq still exhibited high performance (**Table S8; Figure S17**).

**Figure. 5.**
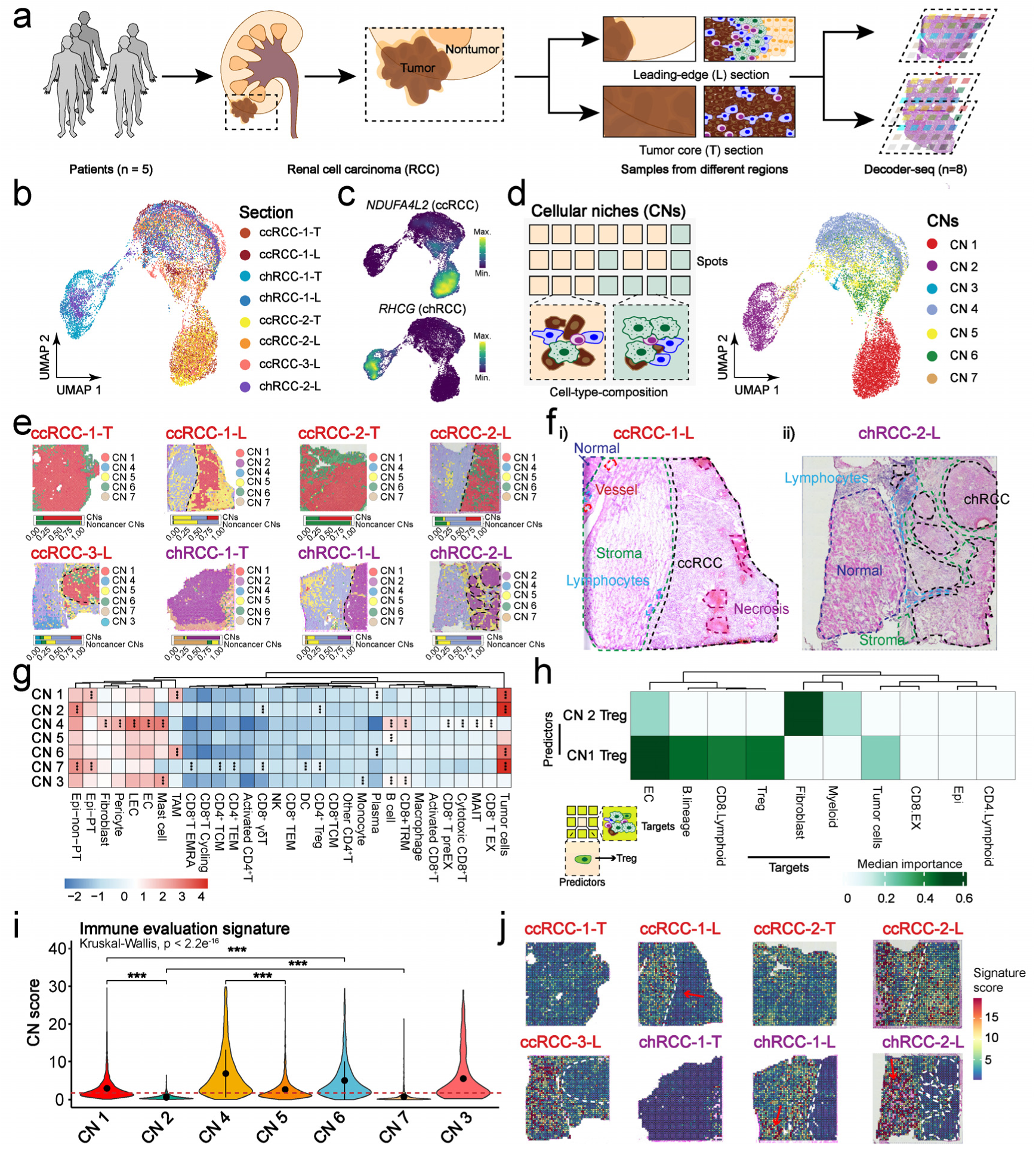
Resolving spatial heterogeneity in RCCs with Decoder-seq. (**a**) Workflow of RCC sample collection and processing for Decoder-seq. **(b)** Uniform manifold approximation and projection (UMAP) plot of spots from all sections. The dot color corresponds to the sample source. **(c)** UMAP plot of the marker gene scores from all sections. The shade of the color represents the scores of the corresponding genes. *NDUFA4L2* was the marker gene for ccRCC, and RHCG was the marker gene for chRCC. **(d)** Schematic of CN definition (left) and UMAP of CNs based on cell-type compositions (right). **(e)** Spatial projection (top) and constituent ratio of CNs (bottom) in each tissue slide. The tumor region is located on the right side of the dotted line. **(f)** H&E images of representative ccRCC and chRCC tissue sections. Pathological annotation of morphological regions into distinct categories including normal renal (dark blue), stromal (green), lymphocyte aggregates (light blue), blood vessel (red), necrosis (purple), and renal cell carcinoma (black). **(g)** Scaled mean cell-type composition within each CN. Asterisks indicate the increased abundance of a specific cell type in a given CN compared with other CNs (two-sided Wilcoxon rank sum test, adjusted (adj.) p < 0.05). Epi-non-PT: Epithelial-non-proximal tubule cell; Epi-PT: Epithelial-proximal tubule cell; LEC: Lymphatic endothelial cell; EC: endothelial cell; TAM: tumor-associated macrophage; CD8^+^ T EMRA: CD8^+^ terminally differentiated effector memory (T EMRA) cell; CD8^+^/CD4^+^ TCM: CD8^+^/CD4^+^ T central memory cell; CD8^+^/CD4^+^ TEM: CD8^+^/CD4^+^ T effector memory cell; NK: natural killer cell; DC: dendritic cell; CD8^+^ TRM: CD8^+^ tissue-resident memory cell; CD8^+^ T EX: Exhausted CD8+ T cell; CD8^+^ T preEX: Precursor exhausted CD8^+^ T cell. MAIT: Mucosal-associated invariant T cell. **(h)** Median importance of Treg abundance for predicting the abundances of other cell types within a 15-spot radius. **(i)** The ratio of cytotoxic CD8^+^ T cells to Tregs in different CNs. **(j)** Spatial projection of the ratio of cytotoxic CD8^+^ T cells to Tregs in each tissue slide.

To reveal the spatial organization of RCCs, we integrated the Decoder-seq data and conducted unsupervised clustering of all spots (**Figure 5b**). Based on the sample composition and marker genes, we identified specific clusters for ccRCC and chRCC, an insular cluster (only containing spots from ccRCC-3-L), and a shared cluster for both subtypes (**Figure 5b-c, Figure S18a, Table S9**). The shared cluster was further divided into four sub-clusters based on deconvoluted cell-type compositions, which together with other clusters, were redefined as seven cellular niches (CNs) (**Figure 5d**, **Figure S18b-c, Methods**). The CN’s spatial projections aligned with pathological annotations for the tissue sections (**Figure 5e-f**). CN 1 and 2 were mainly located in the tumor regions of ccRCC and chRCC samples, respectively, representing the two subtype-specific cancer niches (**Figure 5e-f**). CN 3 was exclusively observed in the normal region of ccRCC-3-L, and the other CNs (4, 5, 6, and 7) were distributed across all sections (**Figure S18d**). These CNs can therefore be regarded as potential spatial building blocks with distinct functions.

To investigate spatial heterogeneity in RCCs, we compared the CNs based on overrepresented cell types and distinct spatial distributions (**Figure 5g**). The subtype-specific CNs 1 and 2 had high abundances of proximal tubule epithelial cells and distal tubule epithelial cells, respectively, in line with their different origins^40, 41^ (**Figure 5g**). Regulatory T cells (Tregs) were the dominant immune cell population in chRCC compared to ccRCC, consistent with the bulk deconvolution of two subtypes from Cancer Genome Atlas (TCGA), suggesting an immunosuppressive TME in chRCC (**Figure S18e-g**). We further observed distinct spatial interaction patterns of Tregs with other cells in the two CNs (**Figure 5h, Methods**). Tregs in CN 1 exhibited the highest predictability for the abundance of endothelial cells, B lineage, CD8 lymphoid, and tumor cells, suggesting a potential vascular recruitment mode^42^. Besides, spatial projection of angiogenic pathways confirmed that ccRCC had higher intra-tumoral vascularization than chRCC, possibly for nutrient delivery and immune cell recruitment (**Figure S18h**). In CN 2, Tregs were the most predictive of fibroblasts, endothelial cells, and myeloid abundance, indicating fibroblast-mediated Treg enrichment in chRCC^43^. Together, these results revealed significant spatial differences in cell composition, immune-infiltration patterns, and cell-to-cell interactions between the two subtypes.

Describing immune cell infiltration patterns based on spatial information could yield additional insights into tumor invasion^36^. We next investigated the spatial immune heterogeneity across different tissue regions (**Figure 5g**). An evaluation signature was devised by calculating the ratio of cytotoxic CD8^+^ T cells (a tumor immune response indicator) to Tregs (an immunosuppressive indicator) in these CNs^44^ (**Figure 5i**). We observed that a highly immune-infiltrated CN 4, mainly located in stromal + normal regions, shaped the variation patterns of CD8^+^ T cells, fibroblasts, and endothelial cells in regulating antitumor immunity (**Figure 5f and Figure 5i**). CN 5 was widely distributed among distinct tissue regions and only displayed a relatively higher abundance of B cells with relatively low immune infiltration. CN 6 and CN 7 were primarily located around tumor edges; the former was immune-infiltrated with tumor-associated macrophages (TAMs) and plasma cells, whereas the latter contained both CD8^+^ T cells and CD4^+^ T cells but remained immunosuppressive due to its lowest ratio (**Figure 5g and Figure 5i**). Thus, when spatially projecting the evaluation signature, we found some “hot” regions within samples presumed to be cold (chRCC-1-L and chRCC-2-L), and a relatively immunosuppressed tumor region within the immune infiltrated ccRCC (ccRCC-1-L) (**Figure 5j, red arrows**). Collectively, Decoder-seq revealed spatial immune heterogeneity across different tissue regions, indicating that immunosuppression or infiltration was not only related to sample subtypes but also to tissue spatial distributions.

### Decoder-seq deciphers potential tumor invasion behaviors associated with clinical outcomes in RCCs

Understanding the TME in primary-stage tumors can reveal the evolutionary trajectory of premalignancy toward malignancy and establish correlations with clinical outcomes^45^. Among the seven CNs, we observed the distribution of CN 5 spots in both subtypes across distinct regions, including tumor core, invasion margin, and stroma + normal tissues (**Figure 5e-f**, **Figure 6a**). Tumor cells in CN 5 exhibited an outward disseminated diffusion pattern from the tumor core to the invasion margin to normal regions, with a small number of tumor cells observed in the stromal + normal zones (**Figure 6a, right**). This is consistent with the characteristics of tumor cells that display swelling and invasive growth, prompting us to consider whether the evolutionary trajectory of tumor invasion occurred in CN 5, following this diffusion path.

**Figure. 6.**
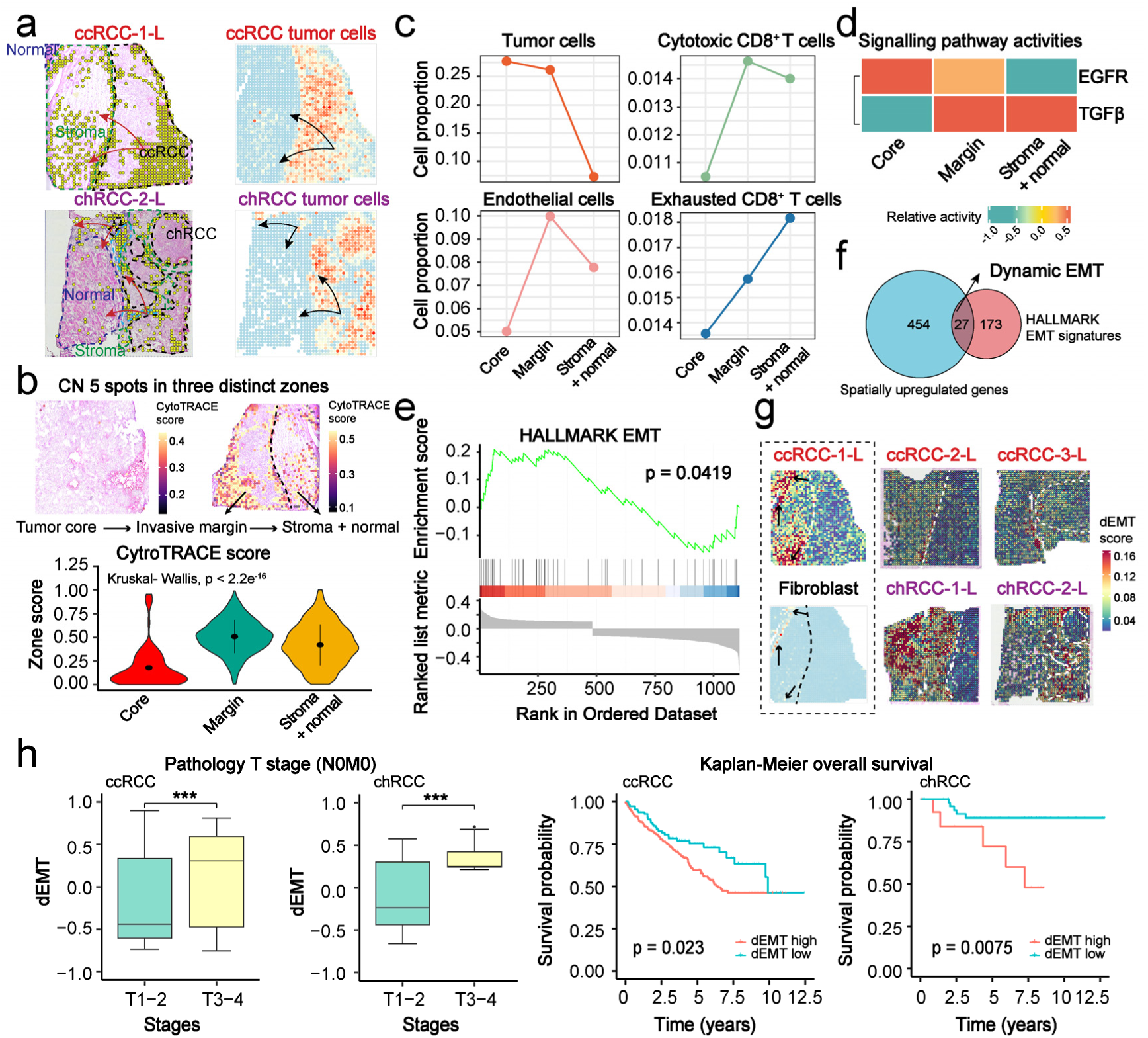
Deciphering the potential spatial evolutionary trajectory of tumor invasion in RCCs. (**a**) Spatial projection of CN 5 (left) and tumor cells (right) in representative tissue sections of ccRCC and chRCC. CN 5 was distributed across distinct tissue regions, and a small number of tumor cells were observed in the stroma + normal CN 5 spots. **(b)** CN 5 spots in representative tissue sections of ccRCC-1-T and ccRCC-1-L were shown their distributions in three distinct tissue zones (top), and violin plots showing the cytroTRACE scores of CN 5 in these zones (bottom). **(c)** The proportion of tumor cells, cytotoxic CD8^+^ T cells, endothelial cells, and exhausted CD8+ T cells in distinct tissue zones. **(d)** Standardized mean PROGENy pathway activities among the spots in CN 5. **(e)** Gene Set Enrichment Analysis (GSEA) of SUSs in the HALLMARKS database of the cancer gene set. **(f)** Venn diagram of the SUSs with the classic EMT signatures in HALLMARKS. **(g)** Spatial projection of dEMT signature on five L sections and spatial distribution of fibroblasts in ccRCC-1-L (black dotted box). **(h)** Survival curves and primary T stages of cases with high and low dEMT scores in TCGA cohorts. Log-rank test was used to measure the statistical significance.

We further employed CytoTRACE^46^ to calculate the transcriptome activity in three zones of CN 5 (**Figure 6b**). Transcriptional activity somewhat correlates with cell proliferation capacity (**Figure S19a-c**). Compared to tumor cores, the invasion margin and stroma + normal zones exhibited significantly higher transcriptome activity, indicating a stronger proliferative potential of cells at tumor edges (**Figure 6b**). Cell compositional analysis revealed an increased number of cytotoxic CD8+ T cells and endothelial cells at the invasion margins (**Figure 6c**). An intense struggle between marginal tumor cells and immune cells may have occurred in this zone, similar to those observed during ecological species invasions. In contrast, the stromal + normal zones exhibited a decrease in cytotoxic CD8^+^ T cells and an increase in exhausted CD8+ T cells (**Figure 6c**). This change may help tumor cells evade immune surveillance, thereby promoting their invasion and proliferation^36^.

We next investigated the spatial variation of molecular signature in CN 5 along the direction of outward disseminated diffusion. PROGENy pathway analysis revealed a down-regulation of the epithelial-related EGFR pathway and an up-regulation of the EMT-related TGFβ pathway in this spatial direction (**Figure 6d**). This may suggest that some tumor cells gradually underwent EMT, acquiring invasive and migratory properties conducive to tumor invasion. To confirm this possibility, we identified a set of genes upregulated along the spatial gradient, referred to as spatially upregulated signatures (SUSs) (**Table S10**). Their functional enrichment analyses revealed the activation of EMT potential in CN 5 (**Figure 6e, Figure S19d, and Table S11**).

EMT is one of the most crucial events in tumor invasion and is associated with a poor prognosis. By intersecting the SUSs with EMT genes in the HALLMARKS database, we identified 27 EMT-related genes, defined as dynamic EMT (dEMT) (**Figure 6f, Table S12**). The spatial projections of the dEMT score on L sections exhibited its extensive distribution in stromal zones of CN 5, suggesting that these regions may serve as the “soil” for tumor invasion (**Figure 6g, Figure S19e**). Notably, we observed the partial dEMT program was activated to colocalize with fibroblasts, consistent with previous studies^47^ (**Figure 6g, black dotted box**). Lastly, we applied the dEMT to the TCGA database and found that advanced RCC patients (stages T3-4) without lymph nodes or distant metastasis had higher dEMT scores than those in stages T1-2 (**Figure 6h**). High dEMT scores were also correlated with an increased likelihood of a poorer prognosis for RCC patients (**Figure 6h**). In contrast, the classic EMT signatures in HALLMARKS failed to predict patients’ prognosis in ccRCC (p = 0.05) (**Figure S19f**). Overall, these findings revealed a potential spatial trajectory of tumor invasion in RCCs and identified EMT-related genes associated with clinical stages and outcomes among RCC patients.

## DISCUSSION

Spatially barcoded array-based spatial transcriptomics technologies have attracted extensive attention for mapping tissue structures and functions with genome-wide coverage. Recently, remarkable methodologies emerged that mainly focused on solving limitations in resolution, improving it from the multicellular level (100 μm) to cellular (10 μm) and even subcellular (∼220 nm) level. Nevertheless, the mRNA capture efficiency and gene detection sensitivity still require optimization to prevent the loss of some important genes due to low expression or partial RNA degradation resulting in low abundance. Moreover, random barcoding strategies necessitate sophisticated and costly position decoding processes, which may limit the affordability and accessibility of such methods to some research communities. Considering that current methods compromise on either the sensitivity, spatial resolution, or cost, we herein develop Decoder-seq by designing a 3D dendrimeric DNA coordinate barcoding array for high-sensitivity spatial transcriptomic analysis with low costs and flexible resolution that can be scaled to near-cellular level.

Decoder-seq demonstrated high sensitivity in gene detection, benefiting from high mRNA capture efficiency on 3D dendrimeric slides. Capturing mRNAs with spatially DNA barcodes is a prerequisite for gene detection in such methods. It requires an adequate modification density of DNA probes on the substrate for efficient collision and contact between the tissue-released mRNAs and probes. The dendrimeric nano-substrate in Decoder-seq offered massive functional groups for densely modifying capture probes, whose density was ∼ an order of magnitude higher than reported spatial transcriptomics methods. Decoder-seq yielded 40.1 UMIs per μm^2^, outperforming existing similar methods. It also exhibited superior sensitivity in detecting lowly-expressed *Olfr*-genes, with an average of 757 genes detected compared to only 132 for the 10× Visium. This advancement may facilitate high-throughput mapping of *Olfr* receptors to glomeruli in the MOB and helps to provide greater insight into olfactory sensory processing.

Decoder-seq also benefitted from high spatial resolution with respect to the deterministic barcoding approach. Deterministic barcoding strategies simplify the fabrication of spatially barcoded arrays and are free of position decoding. Microspotting was commonly used for barcoding, but achieving cellular resolution posed significant technical challenges due to existing instrument limitations. Decoder-seq adopted a microfluidics-assisted coordinate barcoding strategy with exceptional flexibility and precision in spatial resolution (50, 25, and 15 μm). Fan’s group and Peng’s group have utilized this microfluidic barcoding strategy for direct capture and spatial barcoding mRNAs in fixed tissue slides and verified high resolution for spatial sequencing analysis^21, 48^. However, clamping the microchannel chip onto the tissue section requires adding reaction reagents into one hundred inlets for each experiment, which is not user-friendly. In contrast, Decoder-seq’s microfluidics-assisted barcoding array slides are easily mass-fabricated and compatible with 10× Visium’s commercial workflow, making them more easily adopted by researchers for broader applications.

The high cost of current spatial transcriptomics techniques has hindered their accessibility. Decoder-seq can dramatically decrease costs from two aspects. First, the cost of fabricating spatially barcoded arrays is reduced by microfluidics-assisted coordinate barcoding strategy, including a reduced variety of DNA barcodes from combinatorial barcoding, low-concentration DNA required for adopted covalent linkage reaction, and deterministic DNA for decoding-free spatial index. The consumable cost of fabricating a 5 × 5 mm^2^ array is only ∼91 RMB ($0.5/mm^2^; **Table S2-3**), compared favorably to other reported DNA arrays to date. Second, the high gene detection sensitivity allows relatively low sequencing depth for analysis. For instance, to map the MOB (∼ 10.5 mm^2^), Decoder-seq yielded the median number of UMI and genes per spot 40,061 and 8,426 when sequencing ∼299 million reads with a cost of 3,140 RMB (∼$40/mm^2^). In contrast, 10× Visium sequenced ∼399 million reads with only 29,727 and 7,194 per spot, respectively, having a cost of 4,140 RMB (∼$54/mm^2^).

Looking forward, Decoder-seq still has room to further improve performance. First, higher gene detection sensitivity could be achieved by using more favorable RNA capture substrates. For example, G4 dendrimers could be replaced with higher-generation PAMAM dendrimers (G5-G10) with more functional groups to further increase the DNA barcode density. Additionally, regulating the spatial orientation of long single-stranded DNA barcodes may improve mRNA accessibility for capture. Second, it remains a formidable challenge to achieve subcellular resolution using microfluidics-based spatial barcoding methods due to the limitations of soft lithography techniques. This limitation could be overcome by integrating tissue expansion techniques into the Decoder-seq workflow^6^. Third, we here utilized spots and gaps of the same sizes, which resulted in a 25% effective capture area out of the total array area. The gaps between spots could be reduced by decreasing microchannel gaps. Fourth, the current Decoder-seq focused on the polyA-labeled transcriptome. In the future, omics methods such as total transcriptomics, proteomics, and epigenomics could be further combined with it to achieve spatial multi-omics studies^21, 23, 48–50^.

In summary, we developed Decoder-seq that is compatible with high sensitivity, high spatial resolution, and low cost. With 15 μm-spot Decoder-seq, we can readily resolve fine features in the mouse brain and identify lowly-expressed dendrite-enriched genes. Its high sensitivity makes it potentially a valuable tool in clinical research, complementing medical imaging and histopathology data. It was applied to decipher the spatial architecture of highly heterogeneous RCCs and compared the spatial heterogeneity between the ccRCC and chRCC subtypes for the first time. A set of EMT-related genes with a spatial gradient pattern was identified, which strongly correlates with poorer clinical outcomes in RCCs and may thus help predict tumor progression. Decoder-seq as an affordable method for researchers may shine in a wide range of fields such as developmental biology, neuroscience, and clinical research.

## Methods

### Reagents

| REAGENT or RESOURCE | SOURCE | IDENTIFIER |
| --- | --- | --- |
| H<sub>2</sub>SO<sub>4</sub> | Shanghai Lingfeng | 7664-93-9 |
| H<sub>2</sub>O<sub>2</sub> | Shanghai Lingfeng | 7722-84-1 |
| GOPTS | Sigma | 440167 |
| DSS | Thermo Fisher Scientific | 21655 |
| DMSO | Sigma | D2650 |
| PAMAM | Sigma | 412449 |
| Methanol | Shanghai Lingfeng | 67-56-1 |
| 2D commercial slides | Capitalbiotech | 420012 |
| T4 DNA ligase | Vazyme | C301-01 |
| Ethanol | Shanghai Lingfeng | 64-17-5 |
| Methanol | Sigma | 34860 |
| 2-Propanol | Sigma | 19516 |
| Pepsin | Sigma | P7000 |
| Hematoxylin, Mayer's | Dako | S3309 |
| Bluing Buffer | Dako | CS702 |
| Eosin | Sigma | HT110216 |
| Maxima H Minus Reverse Transcriptase | Thermo Fisher Scientific | EP0751 |
| dNTP mix | Thermo Fisher Scientific | R0192 |
| RNase Inhibitor | NEB | M0314L |
| Cy3-dCTP | APExBIO | B8159 |
| dATP | Songon | B110048 |
| dCTP | Songon | B110049 |
| dGTP | Songon | B110050 |
| dTTP | Songon | B110051 |
| Proteinase K | NEB | P8107S |
| Klenow Fragments (3'-5' exo-) | NEB | M0212 |
| Kapa Hotstart HiFi ReadyMix | Roche | KK2602 |
| Kapa qPCR ReadyMix | Roche | KK4601 |
| Equalbit 1 × dsDNA HS Assay Kit | Vazyme | EQ121-01 |
| Ampure XP Beads | Vazyme | N411-03 |
| TruePrep Homo-N7 DNA Library Prep Kit | Vazyme | TD512 |
| Primers, Ligation linkers, DNA barcodes | Songon | See Table S13 |

### Preparation of 3D dendrimeric slides and 2D amino-modified slides

To prepare the 3D dendrimeric slides, glass slides were cleaned and activated by immersion in Piranha solution for 2 h. The slides were then reacted with a solution of glycidyloxypropyltrimethoxysilane (GOPTS) in anhydrous ethanol for 5 h. The slides were washed and dried under an N_2_ stream. After baking in a drying oven for 1 h at 110 °C, the 2D epoxy-modified glass slides were successfully prepared. These slides were then incubated with PAMAM G4 dendrimers in methanol overnight with gentle shaking. Finally, the slides were washed and dried under an N_2_ stream. The PAMAM-modified slides (3D dendrimeric slides) were stored at 4 °C for long-term use.

The 2D amino-modified slides were prepared using a method similar to the one described above for preparing 2D epoxy-modified glass slides. Briefly, the glass slides were cleaned and activated with Piranha solution for 2 h, then treated with a 3-aminopropyltriethoxysilane (APTES) solution in anhydrous ethanol for another 2 h. After sonication in anhydrous ethanol three times as described above and drying under an N_2_ stream, the slides were baked at 110°C for 1 h. Finally, the 2D amino-modified slides were stored at 4 °C.

### Microfluidic device design and fabrication

Briefly, microfluidic devices were designed and fabricated as described for chips of DBiT-seq with certain modifications^21^. Five types of microfluidic chips were here designed with different widths and numbers of channels: 50 channels*50 μm, 50 channels*25 μm, 50 channels*15 μm, 100 channels*25 μm, and 100 channels*15 μm (**Figure S5**). Chrome photomasks and acrylic clamps were ordered from CChip Scientific Instrument Co., Ltd. (Suzhou, China). A thin layer of SU-8 photoresist was spin-coated and patterned with ultraviolet treatment on a silicon wafer. Subsequently, these microfluidic chips were fabricated by soft lithography using polydimethylsiloxane (PDMS). To fabricate PDMS chips, curing reagent A and PDMS base B were mixed in a 1:10 ratio, poured into the patterned SU-8 mold after stirring and degassing, and cured at 120 °C for 20 min. The solidified PDMS chips were cut and peeled off the SU-8 mold. Holes were punched with a 1-mm metal punch pen to form inlets and outlets.

### Generation of surfaces with spatial DNA barcodes

We immobilized spatial DNA barcodes on the surfaces of the 3D dendrimeric slides with two-step reactions. In the first step, DSS was used to crosslink amino-modified DNA barcode X with 3D dendrimeric slides through rapid reactions between the amino and NHS groups. The second step involved the ligation of DNA barcodes X and Y using a complementary linker with T4 DNA ligase, followed by the removal of linkers and enzymes with KOH (**Figure S1b**).

To prevent liquid leakage, we designed a customized rubber hybridization cassette, which contained either one or four separate wells for on-slide reactions, that bound firmly to each slide. Next, 5 μM of DNA barcode X and freshly-prepared DSS in dimethyl sulfoxide (DMSO) were added to 1× PBS and vortexed. The resulting mixture was added to the reaction well of the slide. After incubating for 40 min at room temperature, the solution was removed, and the slide was washed three times. The slides were rinsed with H_2_O and dried under an N_2_ stream.

Due to the high density of amino groups on the 3D dendrimeric slides, excessive unreacted amino groups could have strongly electrostatically absorbed DNA barcode Y, negatively affecting the DNA ligation reaction. The DNA-modified slides were therefore treated with blocking buffer to inactivate any remaining free amino groups. After blocking, the slides were washed and dried under an N_2_ stream. The hybridization cassette was then reinstalled in its original position on the slide for subsequent reactions.

Before the ligation reaction, 5 μM of DNA barcode Y was annealed with 5 μM complementary ligation linker by heating at 95 °C for 2 min, then cooling to 20 °C at a rate of –0.1 °C/s. T4 ligase (40 U/μL) and DNA barcode Y, annealed with a complementary ligation linker, were ligated with DNA barcode X in the reaction well. After incubation at 37 °C for 30 min, the solution was removed, and the slide was continuously washed with wash buffer 1 and H_2_O. After drying, 80 mM KOH was added for 5 min to remove the linker. Slides were rinsed with H_2_O, dried under N_2_, then packaged and stored at 4 °C for future use in investigating tissue permeabilization time.

### AFM characterization

AFM images were obtained with an SPM Multimode and Nanoscope V controller (Bruker, Billerica, MA, USA). Samples were imaged under a ScanAsyst in air mode with Scanasyst air (resonant frequency, f0 = 70 kHz; spring constant, k = 0.4 *N*/*m*) (Bruker).

### Calculation of spatial DNA barcode density on slides

Spatial DNA barcode density on the 2D amino-modified slides and the 3D dendrimeric slides were calculated with a fluorescence spectrophotometer. There were three experimental groups and three control groups, both containing spatial DNA barcodes (oligo-dT primers) labeled with 488 fluorophores. The control groups were used to determine the number of molecules adsorbed to the slides and the experimental groups were used to calculate the number of spatial DNA barcodes. The modification steps for the spatial DNA barcodes used in the experimental groups are described in Methods: Generation of surfaces with spatial DNA barcodes. In the control groups, DNA barcode Y was only hybridized with DNA barcode X without T4 ligase. To perform fluorescence spectrophotometer characterization, we first plotted a standard fluorescence curve using a concentration gradient (0.25, 0.5, 1, 1.5, and 2.5 μM) of DNA barcode Y labeled with the fluorophore 488. We then washed the reaction wells on the slides with 2× SSC three times and recycled the solution. KOH (80 mM) was added to the reaction wells for 5 min to recover free DNA barcode Y. Finally, KOH was mixed with wash buffer to a total volume of 150 μL, which was utilized for fluorescence spectrophotometry analysis.

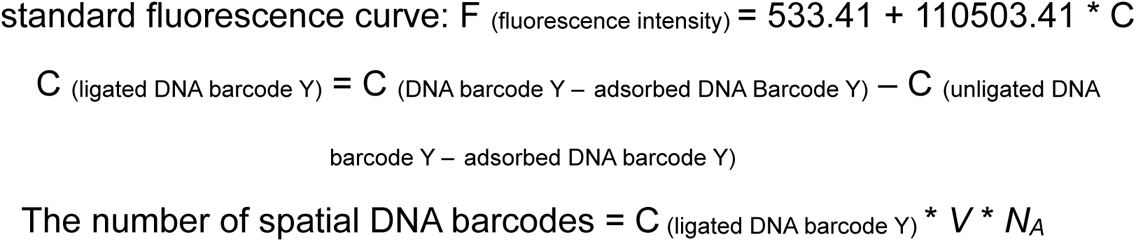

### Generation of dendrimeric DNA coordinate barcoding arrays for Decoder-seq

Dendrimeric DNA coordinate barcoding arrays were generated with microfluidic chips using a method similar to the one described in Methods: Generation of surfaces with spatial DNA barcodes. Two microfluidic chips with orthogonal microfluidic channels in the center were sequentially attached to the 3D dendrimeric slides and firmly held in place with a customized clamp. First, microfluidic chip X (with parallel microchannels) was attached to the 3D dendrimeric slide, assisted by the clamp. The sample comprised a mixture of DNA barcode X and DSS in 1× PBS; 1 µL of the sample was added to the inlets, with each inlet having one unique DNA barcode X. Negative pressure was applied to the outlets with a vacuum pump, introducing samples into the microchannels and causing them to react with the amino groups on the slides.

After incubation for 40 min, the samples were aspirated and the channels were rinsed with a continuous flow of 1 ml of 2× SSC buffer. The clamp and PDMS chip were then removed, and the slide was thoroughly washed with wash buffer 1 and H_2_O.

After the slide dried, the blocking buffer was used to eliminate the remaining unreacted amino groups. A ligation mix of DNA barcode Y (5 μM), ligation linker (5 μM), 1× ligase buffer, and H_2_O was prepared and annealed using a preset program. After annealing, T4 ligase (40 U/μL) was added to each ligation mix. Next, microfluidic chip Y (with channels perpendicular to those of chip X in the barcoding region) was carefully aligned and affixed onto the slide. As described for the DNA barcode X samples, 1 μL of each ligation sample was introduced into the microchannels under negative pressure at the outlets. The ligation reaction was carried out at 37 °C for 30 minutes. Samples were then aspirated and the channels were rinsed with 1 ml of 2× SSC buffer. The slide was disassembled and washed with wash buffer 1 and H_2_O. Linkers were eliminated by treating the sample with 80 mM KOH for 5 min. Finally, the slide was rinsed with H_2_O, dried under a stream of N_2_, and stored at 4 °C. DNA sequences utilized here are shown in **Table S13.**

### Collection and preparation of MOBs and hippocampus

All animal experiments described in this study were conducted in compliance with ethical regulations and approved by the Ethics Committee for the Welfare of Laboratory Animals (license number ACU23-0001).

Male mice (6-8 weeks, Shanghai JieSiJie Laboratory Animals Co.) were euthanized and the whole mouse brain was immediately isolated. The brains were then quickly washed with cold PBS, dried with absorbent paper, and transected along the coronal plane using a sterile surgical blade to divide it into two distinct regions: the MOB and the main brain. These tissues were immediately embedded in cold OCT in plastic molds, frozen on dry ice, and stored at –80 °C before cryosectioning. The tissues were sliced into 10-µm sections with a cryostat at –20 °C. These sections then adhered to the dendrimeric DNA coordinate barcoding arrays for Decoder-seq.

### Collection and preparation of RCC biopsy

The experimental procedures involving human biopsy samples were conducted with the approval of the Ethics Committee of Renji Hospital Affiliated to Shanghai Jiao Tong University School of Medicine (approval number KY2023-013-B). Each donor provided written informed consent before participation.

Eight biopsies from five patients were collected and snap-frozen on dry ice. To determine the RNA quality for analysis, all tissues were assessed using the Qsep 100 Advance (BiOptic Inc.) to assess the degree of RNA degradation. Each tissue was sliced into sections of 10 µm in thickness using a cryostat at – 20 °C. Samples were then trimmed to a suitable size for mounting onto the dendrimeric DNA coordinate barcoding arrays.

### H&E staining and imaging

Tissue sections were incubated at 37 °C for 1 min, then fixed in cool methanol at –20 °C for 30 min. Isopropanol was added to cover each section and the samples were incubated for 1 min. After isopropanol was removed, the sections were air-dried, then sequentially stained with hematoxylin for 7 min, bluing buffer for 2 min, and eosin for 1 min. Sections were incubated at 37 °C for 5 min, then imaged with the Slide Scanning Platform under the bright field.

### Permeabilization and RT

To optimize permeabilization time, a permeabilization experiment was performed. Briefly, these sections were permeabilized with 0.1% pepsin for 6, 12, 18, or 24 min at 37 °C, then washed with 0.1× SSC buffer. mRNAs released from the permeabilized section were captured by the spatial DNA barcodes immobilized on the slide. RT mix was added to synthesize fluorescently labeled cDNA. RT was conducted at 42 °C for 2 h. The sections were then washed three times with 0.1× SSC buffer and digested with tissue digestion buffer (100 mM Tris (pH 8.0), 100 mM NaCl, 2% SDS, 5 mM EDTA, and 16 U/mL

Proteinase K) for 30 min at 37 °C. After tissue removal, the fluorescently-labeled cDNA images were visualized under a fluorescent microscope. Finally, the permeabilization time that yielded the highest fluorescent signal with minimal signal diffusion was selected.

For spatial transcriptomic analysis, the sections were stained with H&E to visualize tissue morphology. Once staining was complete, the sections were used in the Decoder-seq workflow, with permeabilization and RT steps carried out using similar methods as described above.

### Secondary strand synthesis

After RT, tissue sections were washed with 0.1× SSC buffer three times and incubated with 80 mM KOH for 5 min to eliminate all mRNAs from the arrays. Following KOH treatment and washing with EB buffer, second-strand cDNA was synthesized. After synthesis, the arrays were washed with EB buffer three times. Second-strand cDNA was eluted from the arrays via treatment with 35 μL of 80 mM KOH for 10 min, followed by neutralization with 5 μL of 1 M Tris (pH 7.0).

### Amplification, library construction, and sequencing

The second-strand cDNA from each sample was amplified with 50 µL 1× KAPA HiFi Hotstart Ready Mix, 10 μL cDNA primers (8 μM), and 1 μL H_2_O. The following PCR program was used: 98 °C for 3 min; 5 cycles of 98 °C for 20 s, 67 °C for 20 s, and 72 °C for 1 min; n cycles of 98 °C for 20 s, 63 °C for 30 s, and 72 °C for 1 min; final incubation at 72 °C for 5 min. PCR products were purified using VAHTS DNA Clean Beads at a 0.6× bead/sample ratio. The final concentrations were quantified with a Qubit™ dsDNA Assay Kit.

For sequencing library construction, a total of 5 ng of DNA from each sample was prepared with the TruePrep Homo-N7 DNA Library Prep Kit following the manufacturer’s instructions. The resulting libraries were sequenced on an Illumina NovaSeq 6000 in PE150 mode.

### Lateral diffusion analysis

We evaluated the lateral diffusion of Decoder-seq by comparing a nucleus-stained image and a Cy3-dCTP-labeled cDNA image obtained from the same tissue section. Calculations were conducted with ImageJ software in five separate regions to determine the average diffusion distance (**Figure S6**).

### Gene expression matrix generation

To obtain a spatial gene expression matrix, a pipeline was prepared to process the sequencing data. For read 1, only barcodes X and Y with a Levenstein distance of 1 were recovered. Other reads were discarded, as were those with poor UMI quality. Read 2 was aligned to the genome with STAR^51^ and annotation tags were added based on the Drop-Seq pipeline (https://github.com/broadinstitute/Drop-seq/). Finally, a gene expression matrix was constructed for the UMIs and reads per gene associated with each barcode *X_i_Y_j_* for downstream analyses.

### Data processing and clustering analysis

H&E images were used to accurately distinguish between tissue and the background. In most cases, spots outside the target tissue were removed, as were those with fewer than 50 observed genes (UMI > 2). However, the *Olfr* gene counts and spatial maps shown in **Figure 2h** were generated using raw UMI counts without filtration. Genes with counts less than< 10 spatial spots were discarded. Mitochondrial and ribosomal genes were removed. Giotto suite^52^ was used to perform spatial gene expression normalization, plotting, dimensionality reduction, spatial domain detection, and differential gene expression analysis. To incorporate spatial information into the clustering analysis, we utilized a hidden Markov random field (HMRF) model integrated with Giotto and allocated each cell to one of *k* spatial domains based on the accepted beta parameter. Furthermore, the Giotto function “findScanMarkers” was employed to annotate each spatial domain using the identified markers.

### Comparison of Decoder-seq with published methods

MOBs were sequenced with the Decoder-seq and the results were compared to those obtained with previously-published methods. The 10× Visium data used here (**Figure 2d-e**; **Figure S10e-f; Figure 2h**) were sequenced by Shanghai OE Biotech Co. (Shanghai, China). The other two 10× Visium datasets (**Figure 2h**) used to compare the number and type of *Olfr* genes identified were obtained from the 10× Genomics website (https://www.10xgenomics.com/resources/datasets/adult-mouse-olfactory-bulb-1-standard-1) and NCBI (GSE153859). DBiT data were also obtained from NCBI (GSE137986) and Stereo-seq data were downloaded from the China National GeneBank DataBase (https://db.cngb.org/stomics/mosta/download/). The ISS data were generated by Xiamen SEERNA Co. (Xiamen, China).

### Hippocampus analysis

Cell type deconvolution and cell number predictions were performed with CytoSPACE. scRNA-seq data of the mouse hippocampus were obtained from the CARD website (https://yingma0107.github.io/CARD/documentation/05_Experiments.html).

Cell counts were validated using counts obtained from H&E images with the HoVer-Net algorithm (https://arxiv.org/abs/1812.06499). For dendritic enrichment analysis, we used raw UMI counts without filtration and followed the methodology described in Slide-seqV2^12^. To identify differentially expressed genes, we manually selected 58 and 273 spots in the soma and proximal dendrite layer, respectively, for analysis based on their spatial distribution as indicated by the UMI map. Significantly differentially expressed genes were assessed with a two-sample *t*-test.

### RCC data processing and integration

The filtered expression matrix of RCCs was performed in Seurat (v4.3.0) for downstream analysis. The R package “harmony” (v0.1.1) was used to correct batch effects and integrate the expression matrices from different RCC samples^53^.

### Gene set scoring and signaling pathway analysis

The “UCell” package^54^ (v2.2.0) in R was used to score gene sets at the spot level. We used predefined gene sets to estimate cell types^55, 56^. Differential expression analysis was performed using the Wilcoxon test to analyze enrichment scores and gene sets. Unless otherwise stated, those with p < 0.05 were considered statistically significant. For signaling pathway activity analysis, we used the PROGENy model matrix^57^ and analyzed the top 100 genes using the “progeny” package (v1.20.0) in R.

### scRNA-seq references of ccRCC and chRCC

scRNA-seq reference data for ccRCC and chRCC were obtained from public databases^37, 38, 58^. The scRNA-seq data of ccRCC had already undergone filtering and annotation to remove doublets, low-quality cells, and undefined cells. For the two chRCC datasets, we performed standard gene and cell filtering and applied “DoubletFinder” (v2.0.3) to remove doublets. The “FindTransferAnchors” and “TransferData” functions in the Seurat package were used to integrate and annotate the single-cell data from the two chRCC datasets, enabling accurate annotation of non-malignant cell types in chRCC. The annotated cells were then integrated into two datasets: one dataset containing ccRCC tumor cells and all non-tumor cells from both ccRCC and chRCC, and one dataset containing chRCC tumor cells and all non-tumor cells from both ccRCC and chRCC. These two datasets were used in subsequent analyses.

### TCGA data processing

Bulk RNA-seq data for kidney renal clear cell carcinoma (KIRC) and kidney chromophobe (KICH) were downloaded from TCGA (https://portal.gdc.cancer.gov). CIBERSORTx (https://cibersortx.stanford.edu/) was used to perform deconvolution on samples of the two subtypes (ccRCC and chRCC) after down-sampling the single-cell reference data at a ratio of 1:20. This provided cell type proportions for each subtype, which were compared with a Wilcoxon test.

Gene set scoring was conducted on data from TCGA using the R package ‘GSVA’ (v1.46.0)^59^. Optimal cutoff values were determined based on the enrichment scores and survival curves were generated using the R packages “survival” (v3.4.0) and “survminer” (v0.4.9) (**Figure 6h**).

### Cell-type deconvolution and CNs analysis

RCC samples were deconvoluted using “CARD” (v1.0) in R based on ccRCC and chRCC scRNA-seq data. Louvain clustering was performed based on the deconvoluted cell composition results^60^. The overrepresented cell types in each CN were determined using the Wilcoxon test.

### SUSs

To identify SUSs in CN 5, we merged all spots from the eight samples containing this CN. We assigned rank variables based on the positions of the tumor core, margin, and stroma + normal zones. Spearman’s correlation analysis was performed to identify gene sets with correlation coefficients > 0.1 and p-values < 0.05. Outliers with correlation coefficients of 1 were filtered out, resulting in the identification of a gene set with a spatially upregulated pattern specific to CN 5.

The “clusterProfiler” R package (v4.6.2) was used for enrichment analyses of Gene Ontology (GO) terms and 50 HALLMARK tumor pathways in the identified gene set. Gene set enrichment analysis (GSEA) was further performed on the EMT gene set within HALLMARK. dEMTs were classified as the intersection of the SUSs and the classical EMT gene set from HALLMARK using the “Venn” (v1.11) package in R. The gene scores for dEMT were based on UCell.

### Transcriptome activity assessment of CNs

CytoTRACE is a widely used computational framework for single-cell transcriptomics. It is used to assess cellular differentiation potential and stemness by detecting changes in gene expression levels^46^. CytoTRACE may not provide a comprehensive assessment of stemness levels in spatial transcriptomic data, but it can serve as an indicator of RNA abundance. We therefore utilized CytoTRACE (v0.3.3) in R to analyze all spots and defined the scoring as a representation of transcriptome activity in the spots. This definition was supported by the significant positive correlation observed between CytoTRACE scores and the upregulation of KRAS signaling, MYC targets V1, and V2 from the HALLMARK gene sets (**Figure S19b**). We also assessed the correlation between tumor proliferative ability and CytoTRACE scores using cell proliferation-related E2F targets^61^ from the HALLMARK gene sets (**Figure S19c**).

### Spatial interaction analysis

The R package “mistyR” (v1.8.0)^60^ was used to analyze the spatial interactions of Tregs with other cells in CN 1 and CN 2. We integrated the cell types identified by CARD deconvolution and conducted spatial interaction analysis in CN 1 and CN 2 using the “run_misty” function. Specifically, we examined Tregs within each spot and their interactions with cells located within a 15-spot radius. The aggregated estimated median importance scores were interpreted as the spatial component dependency under different spatial contexts in the para-view of a 15-spot radius.

## Data availability

All data generated in this study (all Decoder-seq data, 10x Visium data and ISS data) including raw sequence, expression matrix, space-ranger output and image files will be access under the Gene Expression Omnibus (available upon publication).

## Acknowledgments

The authors would like to thank Dr. Donglei Yang (Renji Hospital) and Dr. Shuai Yang (Renji Hospital) for their assistance in AFM characterization. Dr. Zehui Cao (Renji Hospital), Dr. Ding Ding (Renji Hospital), and Dr. Yang Sun (Renji Hospital) for the support of laboratory space. Dr. Ningning Niu (Renji Hospital), Dr. Zili Yu (Wuhan University), Dr. Xian Huang (Renji Hospital) and Dr. Zhixian Yao (Renji Hospital) for the suggestion on the manuscript. Xin Lin (Shanghai Normal University) and Zhaorun Wu (Xiamen University) for their assistance in bioinformatics analysis. Weixiong Shi (Xiamen University), Jianzhou Feng (Yangzhou University), and Shanqing Huang (Xiamen University) for the preparation of microfluidic chips.

We thank the National Natural Science Foundation of China (22293031, 22004083, 21927806, and 82227801), the National Key R&D Program of China (2019YFA0905800), and Innovative research team of high-level local universities in Shanghai (SHSMU-ZLCX20212601), Shanghai Rising-Star Program (23QA1408200) and China Postdoctoral Science Foundation (Grant No. 2021M702167) for their financial support.

## Author information

These authors contributed equally: Jiao Cao, Zhong Zheng, and Di Sun.

### Authors and Affiliations

Institute of Molecular Medicine, Shanghai Key Laboratory for Nucleic Acid Chemistry and Nanomedicine, Renji Hospital, School of Medicine, Shanghai Jiao Tong University, Shanghai, China

Jiao Cao, Zhong Zheng, Di Sun, Xin Chen, Rui Cheng, Tianpeng Lv, Yu An, Jia Song, Lingling Wu, and Chaoyong Yang.

The MOE Key Laboratory of Spectrochemical Analysis & Instrumentation, Department of Chemical Biology, College of Chemistry and Chemical **Engineering, Xiamen University, Xiamen, China** Chaoyong Yang.

Department of Urology, Renji Hospital, Shanghai Jiao Tong University School of Medicine, Shanghai, China

Zhong Zheng and Junhua Zheng

## Contributions

C.Y.Y. conceived the project. C.Y.Y., L.L.W., and J.C. designed the experiments. J.C., Z.Z., and T.P.L. performed the experiments. J.C., L.L.W., J.S., Z.Z., D.S., and R.C. analyzed the data. J.C., X.C., and Y.A. fabricated microfluidic devices. T.P.L. prepared the microfluidic chip. Z.Z. collected the biopsy samples. L.L.W. and J.C. wrote the manuscript. C.Y.Y., W.L.L., J.S., and J.H.Z. revised the manuscript and supervised the project.

## Corresponding author

Correspondence to Junhua Zheng, Jia Song, Lingling Wu, and Chaoyong Yang.

## Supplementary information

Supplementary Figs. 1–19, Tables 1–13, and a supplementary Excel file

## References

1. Palla, G., Fischer, D.S., Regev, A. & Theis, F.J. Spatial components of molecular tissue biology. Nat. Biotechnol. 40, 308–318 (2022).

2. Rao, A., Barkley, D., Franca, G.S. & Yanai, I. Exploring tissue architecture using spatial transcriptomics. Nature 596, 211–220 (2021).

3. Chen, Y. et al. Mapping Gene Expression in the Spatial Dimension. Small Methods 5, e2100722 (2021).

4. Moses, L. & Pachter, L. Museum of spatial transcriptomics. Nat. Methods 19, 534–546 (2022).

5. Chen, K.H., Boettiger, A.N., Moffitt, J.R., Wang, S. & Zhuang, X. RNA imaging. Spatially resolved, highly multiplexed RNA profiling in single cells. Science 348, aaa6090 (2015).

6. Alon, S. et al. Expansion sequencing: Spatially precise in situ transcriptomics in intact biological systems. Science 371, eaax2656 (2021).

7. Tianyi, C., et al. Rapid and Signal Crowdedness-Robust In-Situ Sequencing through Hybrid Block Coding. bioRxiv, 2022.2011.2016.516714 (2022).

8. Moffitt, J.R., Lundberg, E. & Heyn, H. The emerging landscape of spatial profiling technologies. Nat. Rev. Genet. 23, 741–759 (2022).

9. Ståhl, P.L. et al. Visualization and analysis of gene expression in tissue sections by spatial transcriptomics. Science 353, 78–82 (2016).

10. Vickovic, S. et al. High-definition spatial transcriptomics for in situ tissue profiling. Nat. Methods 16, 987–990 (2019).

11. Rodriques, S.G. et al. Slide-seq: A scalable technology for measuring genome-wide expression at high spatial resolution. Science 363, 1463–1467 (2019).

12. Stickels, R.R. et al. Highly sensitive spatial transcriptomics at near-cellular resolution with Slide-seqV2. Nat. Biotechnol. 39, 313–319 (2021).

13. Cho, C.S. et al. Microscopic examination of spatial transcriptome using Seq-Scope. Cell 184, 3559–3572 e3522 (2021).

14. Chen, A. et al. Spatiotemporal transcriptomic atlas of mouse organogenesis using DNA nanoball-patterned arrays. Cell 185, 1777–1792 e1721 (2022).

15. Fu, X. et al. Polony gels enable amplifiable DNA stamping and spatial transcriptomics of chronic pain. Cell 185, 4621–4633 e4617 (2022).

16. Longo, S.K., Guo, M.G., Ji, A.L. & Khavari, P.A. Integrating single-cell and spatial transcriptomics to elucidate intercellular tissue dynamics. Nat. Rev. Genet. 22, 627–644 (2021).

17. Zhu, K.W. et al. Decoding the olfactory map through targeted transcriptomics links murine olfactory receptors to glomeruli. Nat. Commun. 13, 5137 (2022).

18. Wang, I.H. et al. Spatial transcriptomic reconstruction of the mouse olfactory glomerular map suggests principles of odor processing. Nat. Neurosci. 25, 484–492 (2022).

19. Kvastad, L. et al. The spatial RNA integrity number assay for in situ evaluation of transcriptome quality. *Commun*. Biol. 4, 57 (2021).

20. Wu, A.R., Wang, J., Streets, A.M. & Huang, Y. Single-Cell Transcriptional Analysis. Annu. Rev. Anal. Chem. 10, 439–462 (2017).

21. Liu, Y. et al. High-Spatial-Resolution Multi-Omics Sequencing via Deterministic Barcoding in Tissue. Cell 183, 1665–1681 e1618 (2020).

22. Arima, H. & Keiichi, M. Recent Findings Concerning PAMAM Dendrimer Conjugates with Cyclodextrins as Carriers of DNA and RNA. Sensors 9, 6346–6361 (2009).

23. Vickovic, S. et al. SM-Omics is an automated platform for high-throughput spatial multi-omics. Nat. Commun. 13, 795 (2022).

24. Wirth, J. et al. Spatial transcriptomics using multiplexed deterministic barcoding in tissue. Nat. Commun. 14, 1523 (2023).

25. Lein, E.S. et al. Genome-wide atlas of gene expression in the adult mouse brain. Nature 445, 168–176 (2007).

26. Larsson, C., Grundberg, I., Söderberg, O. & Nilsson, M. In situ detection and genotyping of individual mRNA molecules. Nat. Methods 7, 395–397 (2010).

27. Vassar, R. et al. Topographic organization of sensory projections to the olfactory bulb. Cell 79, 981–991 (1994).

28. Yoon, Y.J. et al. Glutamate-induced RNA localization and translation in neurons. Proc. Natl. Acad. Sci. 113, E6877–E6886 (2016).

29. Steward, O. & Worley, P. Local Synthesis of Proteins at Synaptic Sites on Dendrites: Role in Synaptic Plasticity and Memory Consolidation? Neurobiol. Learn. Mem. 78, 508–527 (2002).

30. Kosik, K.S. Life at Low Copy Number: How Dendrites Manage with So Few mRNAs. Neuron 92, 1168–1180 (2016).

31. Vahid, M.R. et al. High-resolution alignment of single-cell and spatial transcriptomes with CytoSPACE. Nat. Biotechnol., 1–6 (2023).

32. Tushev, G. et al. Alternative 3’ UTRs Modify the Localization, Regulatory Potential, Stability, and Plasticity of mRNAs in Neuronal Compartments. Neuron 98, 495–511 e496 (2018).

33. Ainsley, J.A., Drane, L., Jacobs, J., Kittelberger, K.A. & Reijmers, L.G. Functionally diverse dendritic mRNAs rapidly associate with ribosomes following a novel experience. Nat. Commun. 5, 4510 (2014).

34. Nakayama, K. et al. RNG105/caprin1, an RNA granule protein for dendritic mRNA localization, is essential for long-term memory formation. eLife 6, e29677 (2017).

35. Cajigas, I.J. et al. The local transcriptome in the synaptic neuropil revealed by deep sequencing and high-resolution imaging. Neuron 74, 453–466 (2012).

36. Fu, T. et al. Spatial architecture of the immune microenvironment orchestrates tumor immunity and therapeutic response. J. Hematol. Oncol. 14, 98 (2021).

37. Zhang, Y. et al. Single-cell analyses of renal cell cancers reveal insights into tumor microenvironment, cell of origin, and therapy response. Proc. Natl. Acad. Sci. USA 118, e2103240118 (2021).

38. Su, C. et al. Single-Cell RNA Sequencing in Multiple Pathologic Types of Renal Cell Carcinoma Revealed Novel Potential Tumor-Specific Markers. Front. Oncol. 11, 719564 (2021).

39. Kansler, E.R. et al. Cytotoxic innate lymphoid cells sense cancer cell-expressed interleukin-15 to suppress human and murine malignancies. Nat. Immunol. 23, 904–915 (2022).

40. Sanchez, D.J. & Simon, M.C. Genetic and metabolic hallmarks of clear cell renal cell carcinoma. Biochim. Biophys. Acta, Rev. Cancer 1870, 23–31 (2018).

41. Hsieh, J.J., Le, V., Cao, D., Cheng, E.H. & Creighton, C.J. Genomic classifications of renal cell carcinoma: a critical step towards the future application of personalized kidney cancer care with pan-omics precision. J. Pathol. 244, 525–537 (2018).

42. Certo, M. et al. Endothelial cell and T-cell crosstalk: Targeting metabolism as a therapeutic approach in chronic inflammation. Br. J. Pharmacol. 178, 2041–2059 (2021).

43. de Visser, K.E. & Joyce, J.A. The evolving tumor microenvironment: From cancer initiation to metastatic outgrowth. Cancer Cell 41, 374–403 (2023).

44. Kareva, I. Metabolism and Gut Microbiota in Cancer Immunoediting, CD8/Treg Ratios, Immune Cell Homeostasis, and Cancer (Immuno)Therapy: Concise Review. Stem Cells 37, 1273–1280 (2019).

45. Lewis, S.M. et al. Spatial omics and multiplexed imaging to explore cancer biology. Nat. Methods 18, 997–1012 (2021).

46. Gulati, G.S. et al. Single-cell transcriptional diversity is a hallmark of developmental potential. Science 367, 405–411 (2020).

47. Szabo, P.M. et al. Cancer-associated fibroblasts are the main contributors to epithelial-to-mesenchymal signatures in the tumor microenvironment. Sci. Rep. 13, 3051 (2023).

48. Jiang, F. et al. Simultaneous profiling of spatial gene expression and chromatin accessibility during mouse brain development. Nat. Methods, 1–10 (2023).

49. Liu, Y. et al. High-plex protein and whole transcriptome co-mapping at cellular resolution with spatial CITE-seq. Nat. Biotechnol., 1–5 (2023).

50. Ben-Chetrit, N. et al. Integration of whole transcriptome spatial profiling with protein markers. Nat. Biotechnol., 1–6 (2023).

51. Dobin, A. et al. STAR: ultrafast universal RNA-seq aligner. Bioinformatics 29, 15–21 (2013).

52. Dries, R. et al. Giotto: a toolbox for integrative analysis and visualization of spatial expression data. Genome Biol. 22, 78 (2021).

53. Wu, R. et al. Comprehensive analysis of spatial architecture in primary liver cancer. Sci. Adv. 7, eabg3750 (2021).

54. Andreatta, M. & Carmona, S.J. UCell: Robust and scalable single-cell gene signature scoring. Comput. Struct. Biotechnol. J. 19, 3796–3798 (2021).

55. Aran, D., Hu, Z. & Butte, A.J. xCell: digitally portraying the tissue cellular heterogeneity landscape. Genome Biol. 18, 220 (2017).

56. He, Y., Jiang, Z., Chen, C. & Wang, X. Classification of triple-negative breast cancers based on Immunogenomic profiling. J. Exp. Clin. Cancer Res. 37, 327 (2018).

57. Schubert, M. et al. Perturbation-response genes reveal signaling footprints in cancer gene expression. Nat. Commun. 9, 20 (2018).

58. Li, Y. et al. Histopathologic and proteogenomic heterogeneity reveals features of clear cell renal cell carcinoma aggressiveness. Cancer Cell 41, 139–163.e117 (2023).

59. Hänzelmann, S., Castelo, R. & Guinney, J. GSVA: gene set variation analysis for microarray and RNA-seq data. BMC Bioinformatics 14, 7 (2013).

60. Kuppe, C. et al. Spatial multi-omic map of human myocardial infarction. Nature 608, 766–777 (2022).

61. Kent, L.N. & Leone, G. The broken cycle: E2F dysfunction in cancer. Nat. Rev. Cancer 19, 326–338 (2019).

