## Supplementary Information of Supplementary Figures and Tables for "Dendrimeric DNA Coordinate Barcoding Design for Spatial RNA Sequencing"

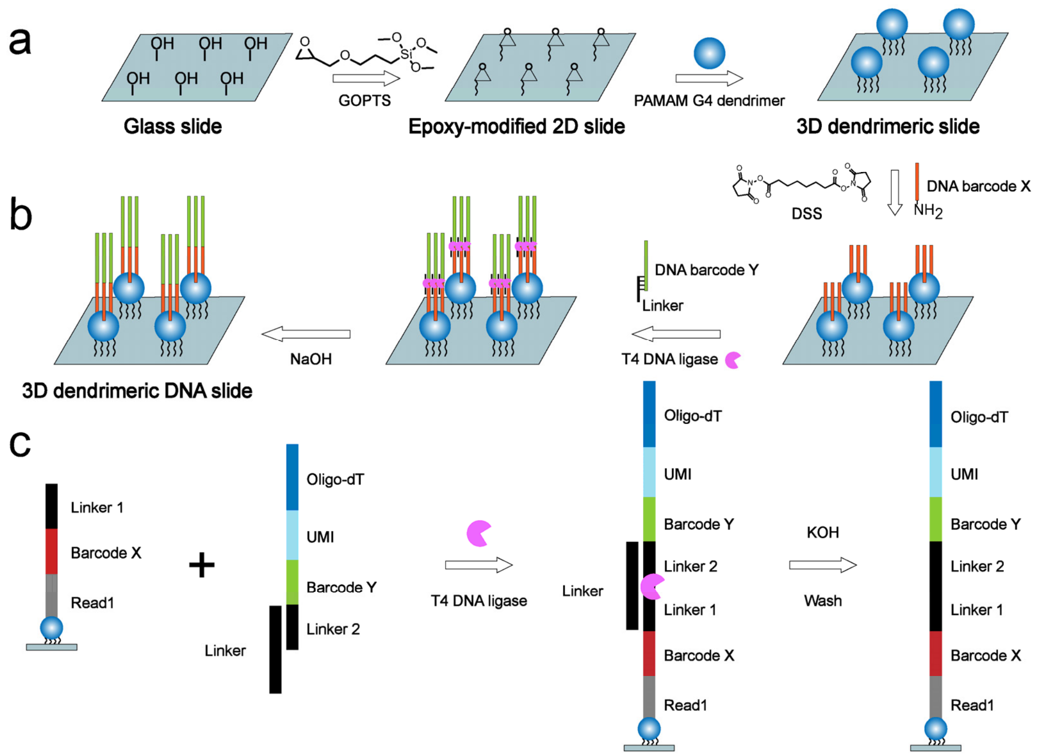

**Supplementary Figure S1. Fabrication and DNA modification of a three-dimensional (3D) dendrimeric slide. (a)** Workflow showing fabrication of a 3D dendrimeric slide. A glass slide activated with hydroxyl groups was reacted with glycidyloxypropyltrimethoxysilane (GOPTS) to form an epoxy-modified two-dimensional (2D) slide. Poly(amidoamine) (PAMAM) G4 dendrimers were then introduced to generate the 3D dendrimeric slide. **(b)** Workflow showing the process by which the spatial DNA barcodes were modified to the slide. Disuccinimidyl suberate (DSS) reagent containing double-terminal N-HydroxySuccinimide (NHS) groups were used to cross-link DNA barcode X to the 3D dendrimeric slide. Through a ligation reaction using T4 DNA ligase, DNA barcode Y (annealed with the linker) was ligated with DNA barcode X. **(c)** Schematic representation of the DNA structure and the ligation reaction used to ligate DNA barcode X and Y on the 3D dendrimeric slide.

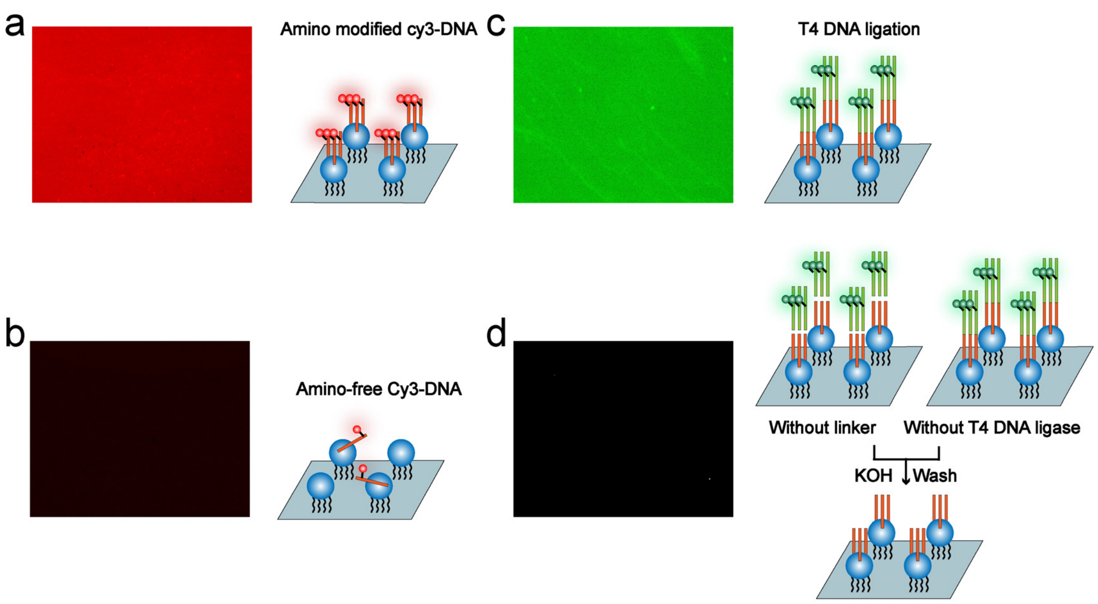

**Supplementary Figure S2. Fluorescence characterization of covalent modification and ligation reactions on 3D dendrimeric slides using DNA probes. (a-b)** Fluorescence images of amino-modified Cy3-DNA probes (**a**) and amino-free Cy3-DNA probes (**b**) on the 3D dendrimeric slide. Fluorescence was clearly observed in the 3D dendrimeric slides after incubation with amino-modified Cy3-DNA, indicating the covalent attachment of amino-modified DNA probes (**a**). Cy3-DNA probes without amino modifications could not be crosslinked and therefore showed no obvious fluorescence on the slide (**b**). **(c-d)** Fluorescence images of Alexa 488-labeled DNA barcode Y on the 3D dendrimeric slide. Alexa 488-labeled DNA barcode Y was ligated to DNA barcode X, which was immobilized on the slide, through a ligation reaction (**c**). The ligation reaction could not occur without a linker DNA or T4 DNA ligase; no fluorescence was observed on the slide in the absence of these components (**d**).

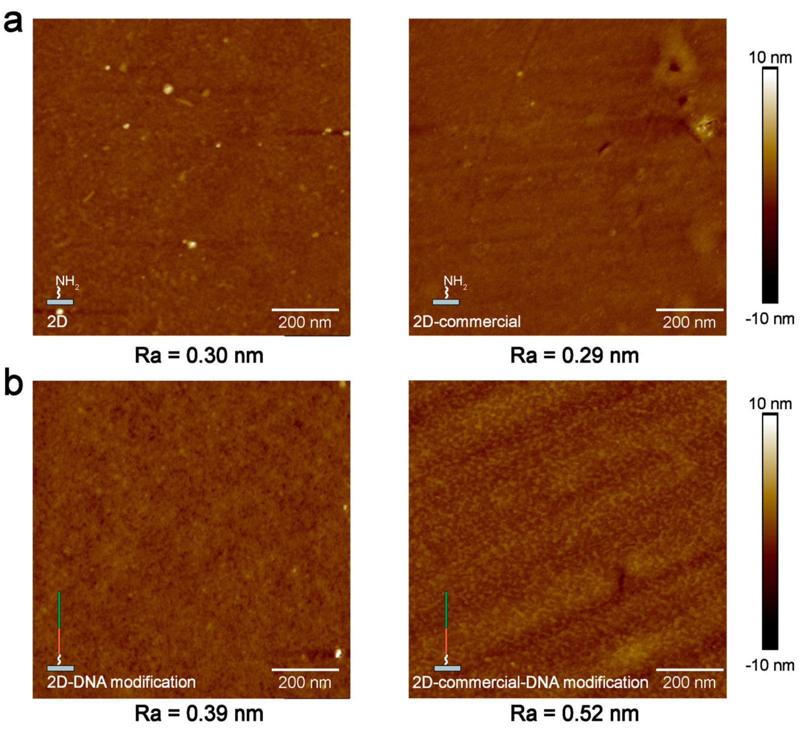

**Supplementary Figure S3. Atomic force microscopic (AFM) imaging for surface morphology characterization of 2D slides. (a)** AFM imaging for surface morphology characterization of 2D amine-modified slide. Left: homemade 2D amine-modified slides; right: commercially-generated 2D amine-modified slides. **(b)** AFM imaging for surface morphology characterization of 2D amine-modified slides (Left: homemade slides; right: commercially-generated slides) modified with spatial DNA barcodes.

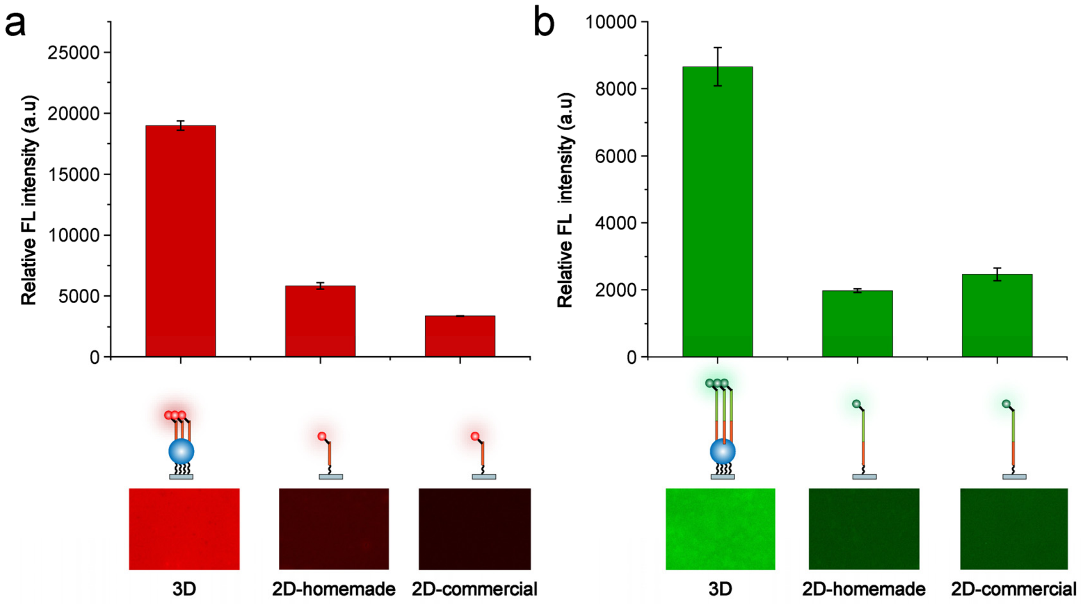

**Supplementary Figure S4. Comparison of fluorescence intensity between 3D dendrimeric slides and 2D planar slides. (a)** Bottom: fluorescence images of a 3D dendrimeric slide and 2D planar slides (2D homemade and 2D commercial) after covalent linkage with Cy3-labeled DNA barcode X. Top: relative fluorescence (FL) intensity as quantified from the images. **(b)** Bottom: fluorescence images of a 3D dendrimeric slide and 2D planar slides after the Alexa 488-labeled DNA barcode Y was ligated with DNA barcode X. Top: relative FL intensity as quantified from the images. Relative FL intensity was calculated as the absolute FL intensity with the background FL intensity subtracted. Data represent the mean ± standard deviation from three independent replicates.

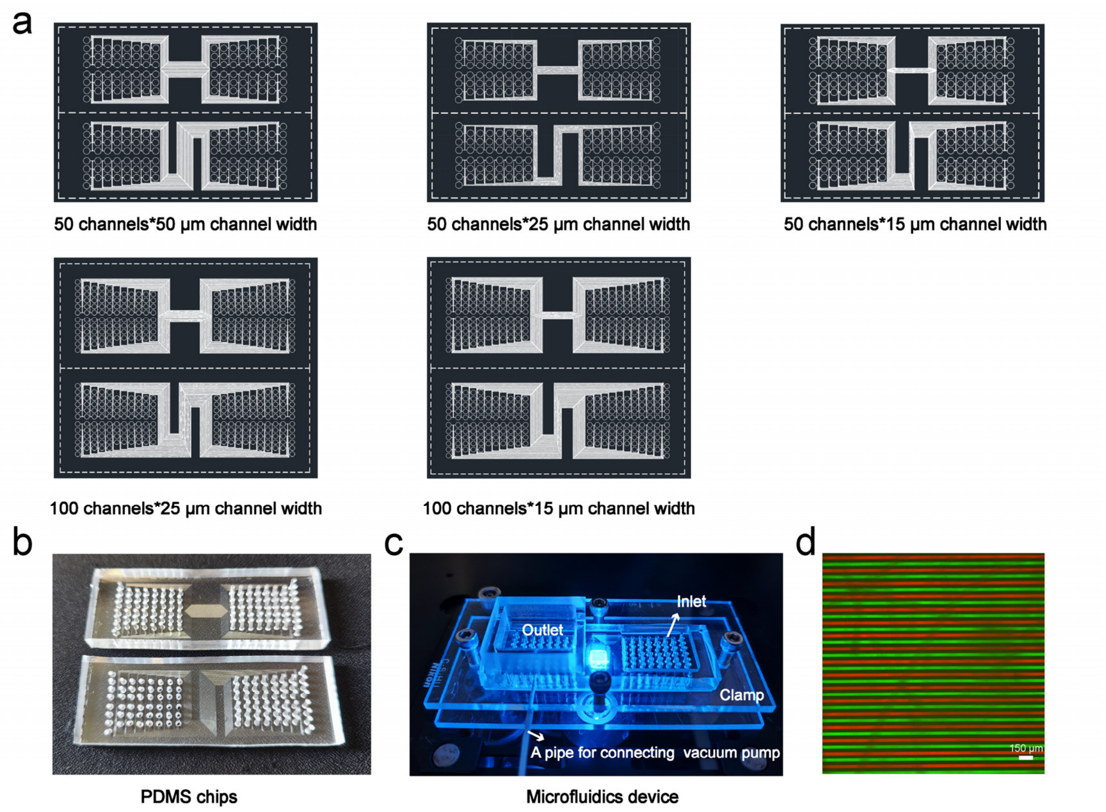

**Supplementary Figure S5. Microfluidic device design. (a)** AutoCAD representations of polydimethylsiloxane (PDMS) chips with varying numbers of channels and widths. Upper left: 50 channels of 50 μm in width; upper middle: 50 channels of 25 μm in width; upper right: 50 channels of 15 μm in width; lower left: 100 channels of 25 μm in width; lower right: 100 channels of 15 μm in width. **(b)** A pair of PDMS chips (with 50 μm channel widths) possessing mutually perpendicular microchannels for spatial barcoding. **(c)** The microfluidic device used for Decoder-seq. A PDMS chip was placed on a slide and clamped using two acrylic plates and screws. The DNA barcode reagents were pipetted into the inlets and drawn into the microchannels through vacuum pressure applied to the roof cap of the outlets, which were situated on the other side of the PDMS chip. **(d)** Measurement of possible crosstalk between neighboring channels by alternately flowing Cy3 (red)-labeled DNA barcode X and Alexa 488 (green)-labeled DNA barcode Y through adjacent channels.

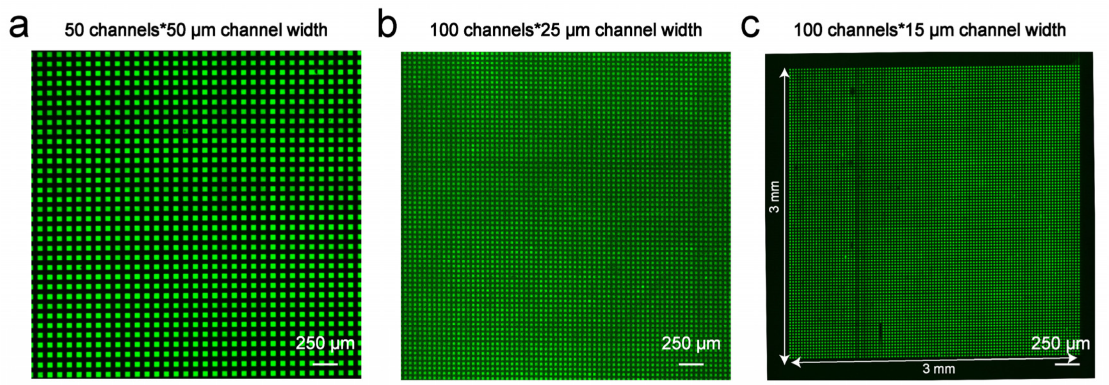

**Supplementary Figure S6. Fluorescence images of 3D dendrimeric DNA coordinate barcoding arrays. (a-c)** Fluorescence images of 3D dendrimeric DNA coordinate barcoding arrays consisting of 2,500 spots of 50 µm in diameter with a total capture area of 25 mm$^2$ **(a),** 10,000 spots of 25 µm in diameter with a total capture area of 25 mm$^2$ **(b)**, and 10,000 spots of 15 µm in diameter with a total capture area of 9 mm$^2$ **(c)**.

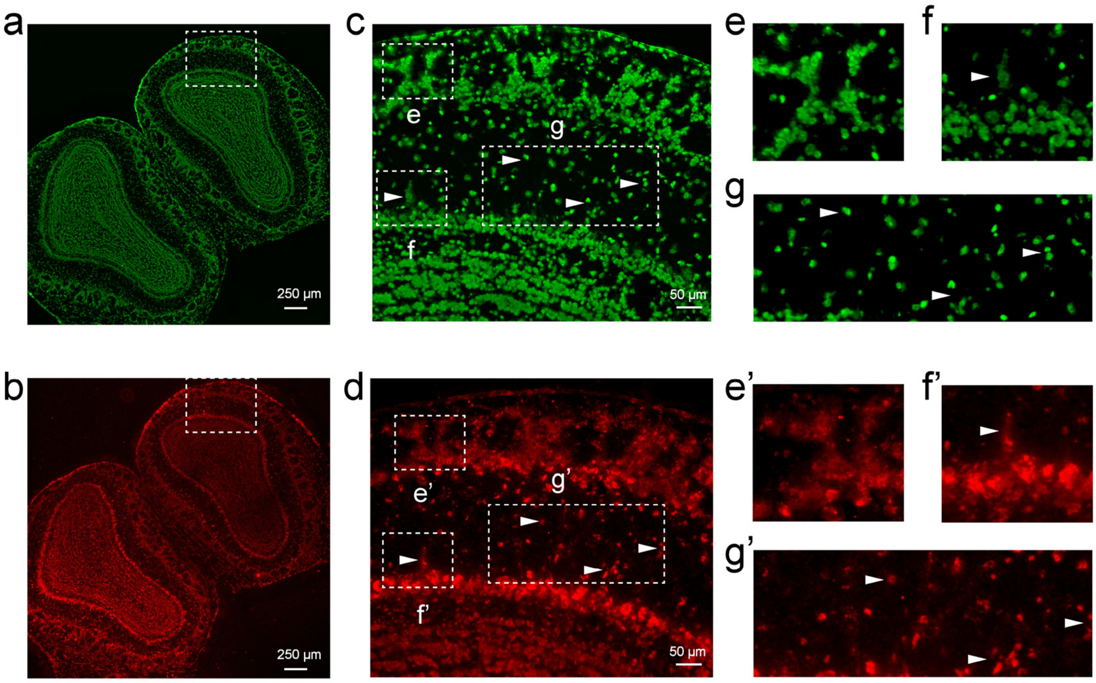

**Supplementary Figure S7. Spatially-localized cDNA synthesis. (a)** Fluorescence image of a mouse olfactory bulb (MOB) tissue section stained with nucleic acid dyes. **(b)** Fluorescence image of cDNA synthesized with Cy3-labeled dCTP after tissue removal. **(c)** Magnified view of the section outlined in white in **a**. **(d)** Magnified view of the section outline in white in **b**. **(e-g)** Magnified views of the sections outlined in white in **c. (e'-g')** Magnified views of the sections outlined in white in **d.** The distribution of fluorescent cDNAs showed a similar pattern to that of the nucleic acid-stained tissue section after tissue removal, indicating that the cDNAs were localized to the areas under the cells and there was negligible diffusion.

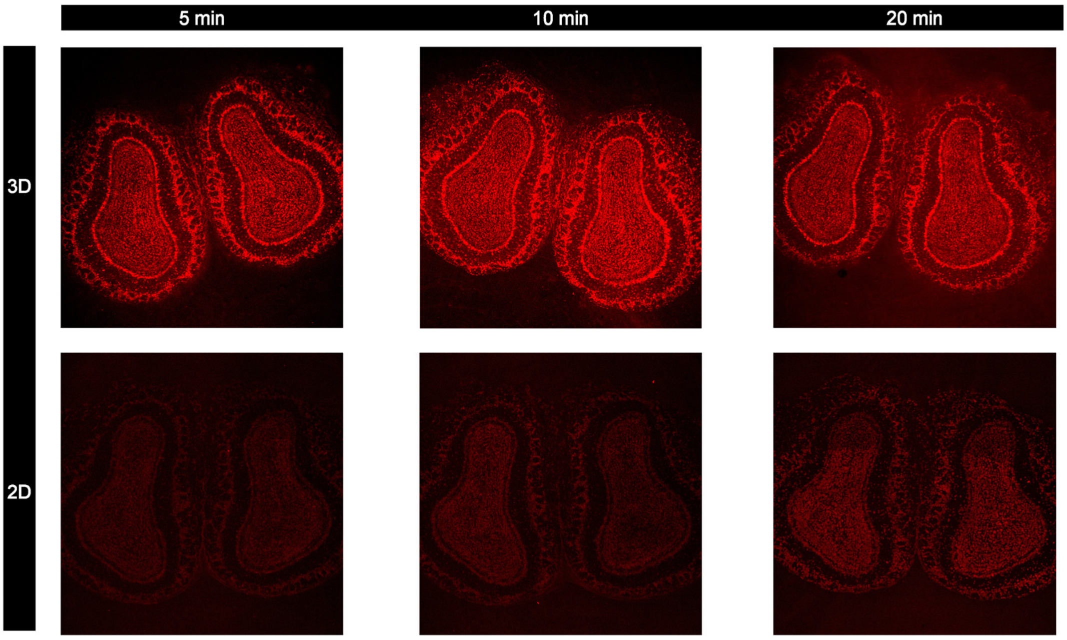

**Supplementary Figure S8. Comparison of fluorescent cDNA patterns from MOB tissues on 3D dendrimeric DNA slides and 2D planar DNA slides. (a-b)** Time series showing fluorescent cDNA footprints on the 3D dendrimeric slide **(a)** and the 2D planar slide **(b)** corresponding to varying pepsin permeabilization incubation times (5, 10, and 20 min). The fluorescent cDNA signals were significantly stronger on the 3D dendrimeric DNA slides than on the 2D planar DNA slides.

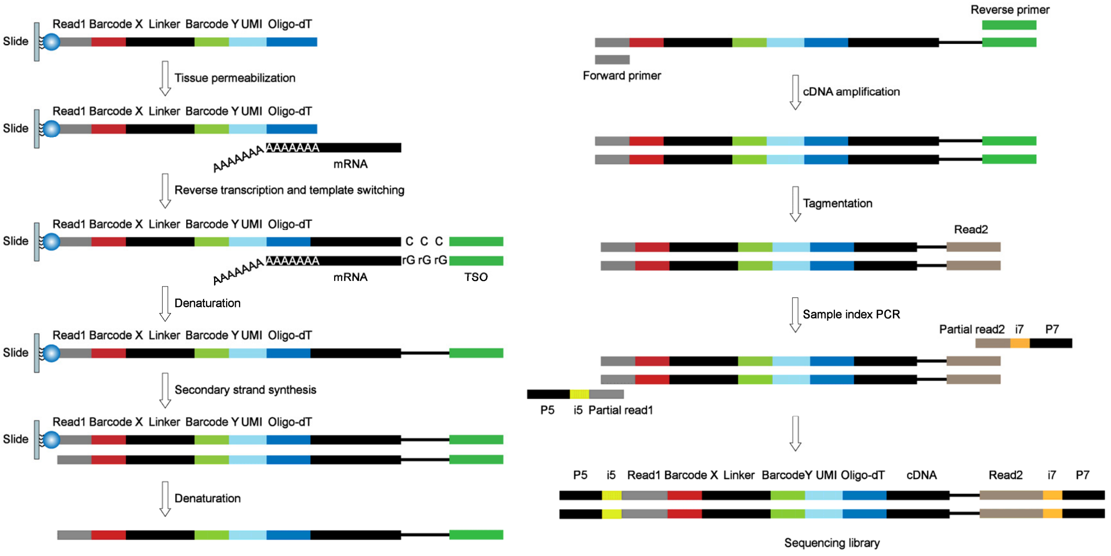

**Supplementary Figure S9. Reverse transcription, second-strand cDNA synthesis, and library construction workflow.** A detailed experimental procedure is described in the **Methods**.

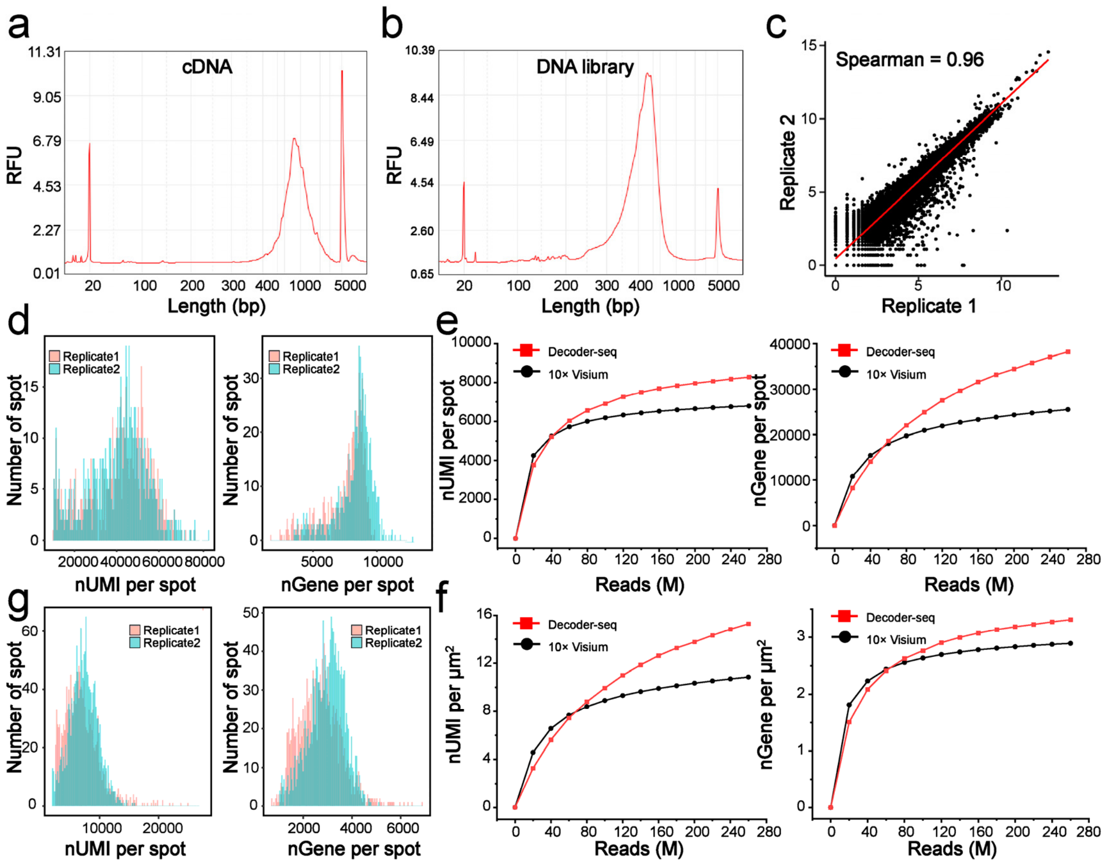

**Supplementary Figure S10. Quality evaluation and comparison of MOB data. (a)** Distribution of cDNA amplicon sizes. **(b)** Distribution of DNA sequencing library sizes. **(c)** Gene consistency of results generated from two replicate MOB sections (Spearman's correlation coefficient = 0.96). **(d)** Distribution of detected unique molecular identifiers (UMIs) (left) and genes (right) per spot from two replicate MOB sections with 50-µm-spot Decoder-seq. **(e, f)** Saturation curves of 50-µm-spot Decoder-seq and 55-µm-spot 10× Visium for median UMI (left) and gene (right) number per spot (**e**) and per µm$^2$ (**f**) versus the raw reads. **(g)** Distribution of detected UMIs (left) and genes (right) from two replicate MOB sections with 15-µm-spot Decoder-seq.

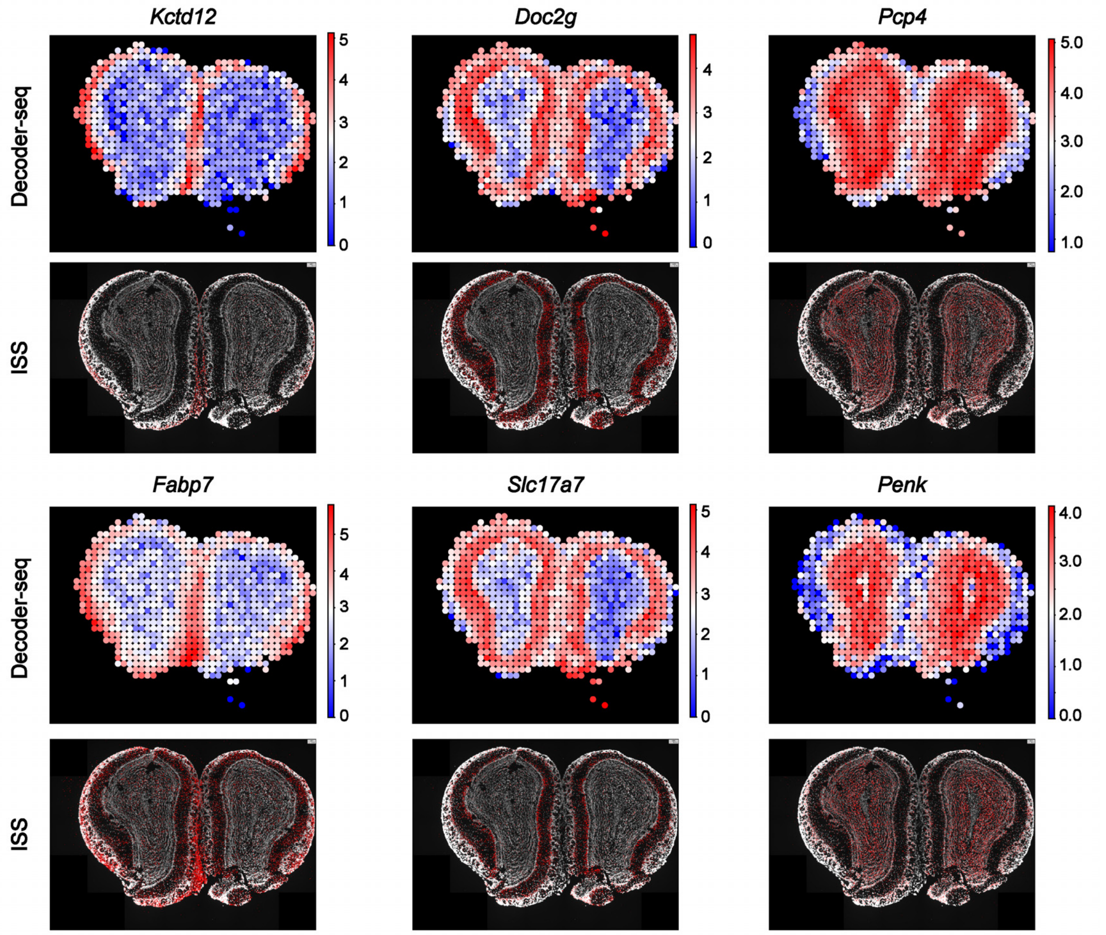

**Supplementary Figure S11. Comparison of Decoder-seq and ISS for selected genes (*Kctd12*, *Doc2g*, *Pcp4*, *Fabp7*, *Slc17a7*, and *Penk*).**

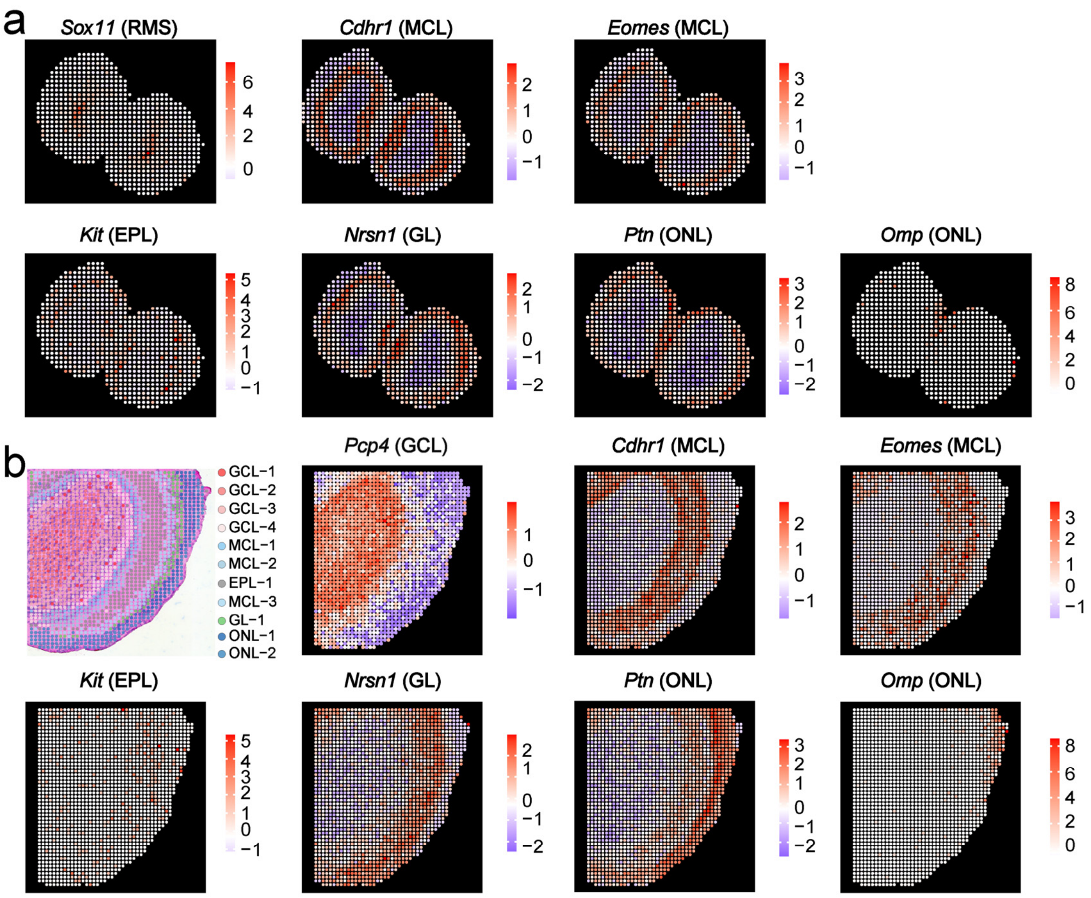

**Supplementary Figure S12. Spatial gene expression patterns of region-specific marker genes in MOBs. (a)** Spatial gene expression patterns of region-specific marker genes in MOBs as determined with 50-µm-spot Decoder-seq. The marker genes and corresponding regions were *Sox11* (RMS), *Cdhr1* and *Eomes* (MCL), *Kit* (EPL), *Nrsn1* (GL), and *Ptn* and *Omp* (ONL). **(b)** Unsupervised clustering of the MOB section and spatial gene expression patterns of region-specific marker genes as determined with 15-µm-spot Decoder-seq. The marker genes and corresponding regions were *Pcp4* (GCL), *Cdhr1* and *Eomes* (MCL), *Kit* (EPL), *Nrsn1* (GL), and *Ptn* and *Omp* (ONL). RMS, rostral migratory stream; GCL, granule cell layer; MCL, mitral and tufted cell layers; EPL, external plexiform layer; GL, glomerular layer; ONL, olfactory neuron layer.

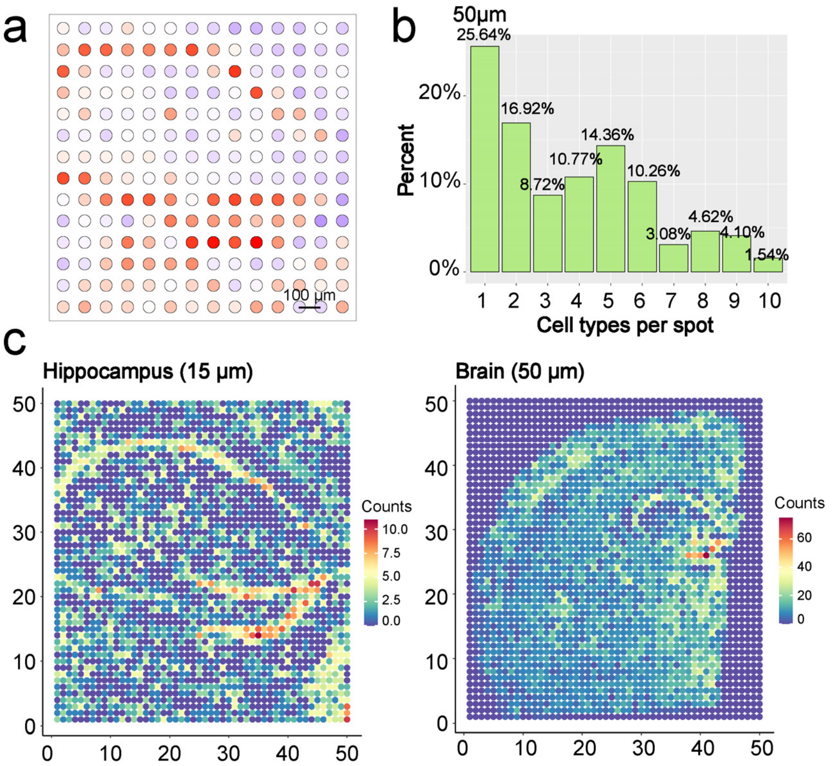

**Supplementary Figure S13. Comparison of mouse hippocampus data from 50-µm-spot and 15-µm-spot Decoder-seq. (a)** Spatial distribution of UMIs in the hippocampus with 50-µm-spot Decoder-seq. **(b)** Number of cell types assigned to each 50-µm spot. **(c)** Heat maps showing cell numbers in each15-µm-spot (left) and 50-µm-spot (right) of Decoder-seq, as determined by H&E based-cell segmentations on the mouse hippocampus (left) and the brain (right).

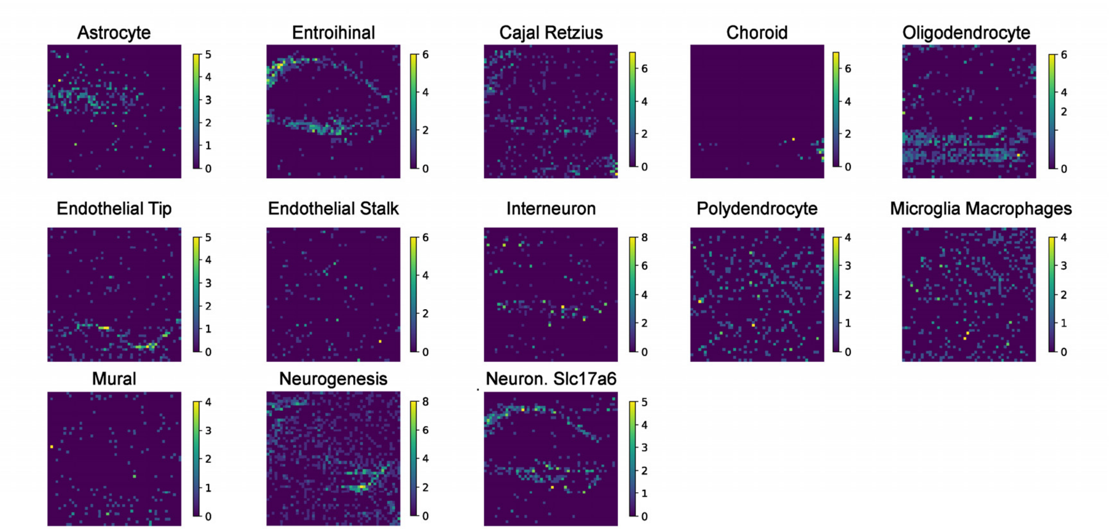

**Supplementary Figure S14. Heat maps showing classical neuronal and non-neuronal cell types in the hippocampus as detected with 15-µm-spot Decoder-seq.**

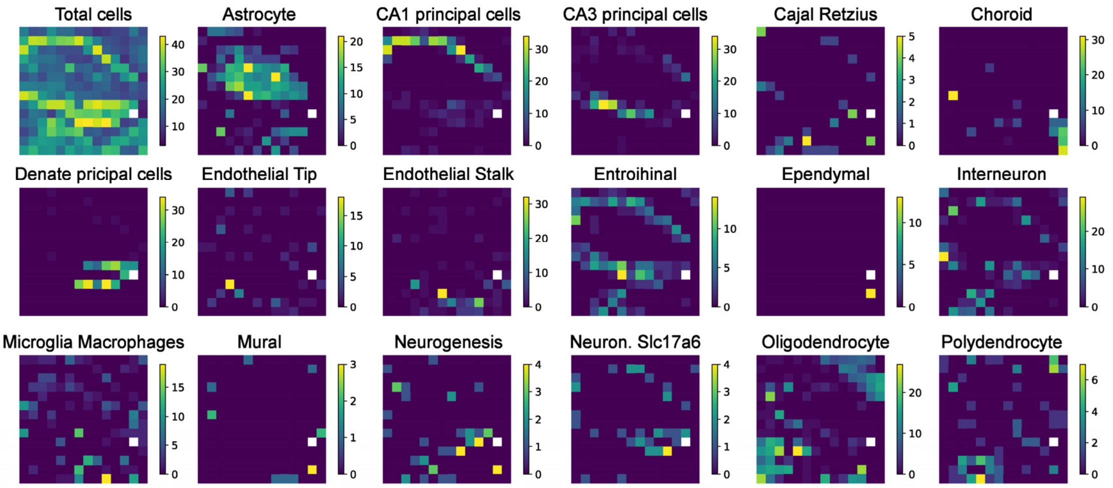

**Supplementary Figure S15. Heat maps showing classical neuronal and non-neuronal cell types in the hippocampus as detected with 50-μm-spot Decoder-seq.**

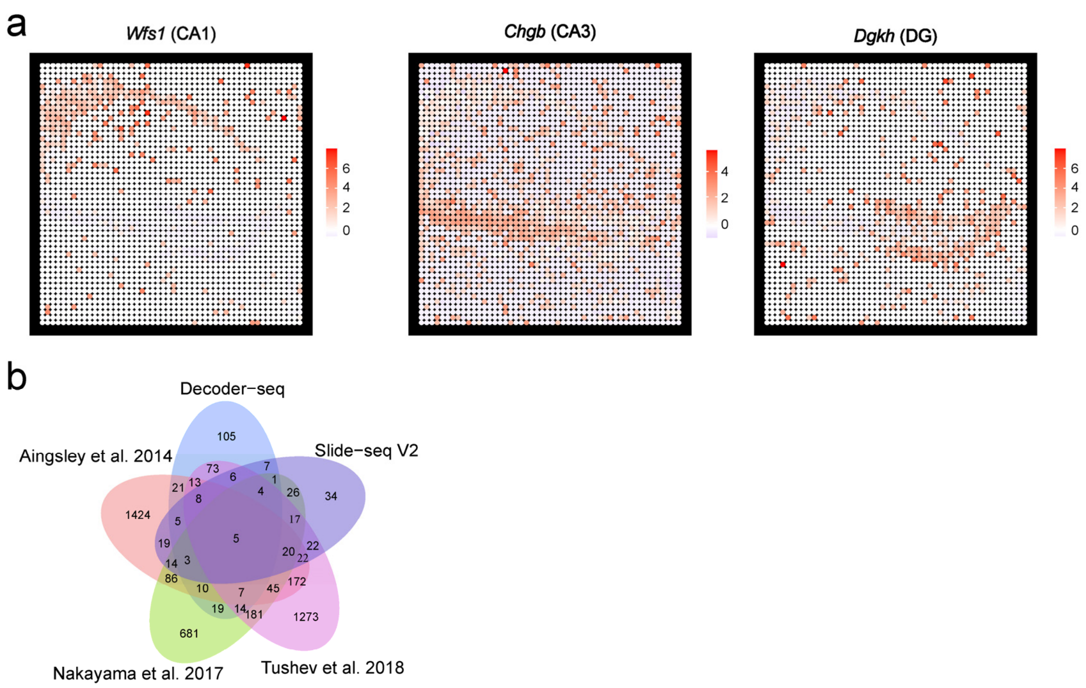

**Supplementary Figure S16. Spatial gene expression of region-specific marker genes in the mouse hippocampus as determined with Decoder-seq. Related to Figure 4. (a)** Left: *Wfs1* for CA1; middle: *Chgb* for CA3; right: *Dgkh* for dentate gyrus (DG). **(b)** Overlap between dendrite-enriched genes identified in the present study and by Aingsley et al., 2014[1], Tushev et al., 2018[2], Nakayama et al., 2017[3], and Slide-seqV2[4].

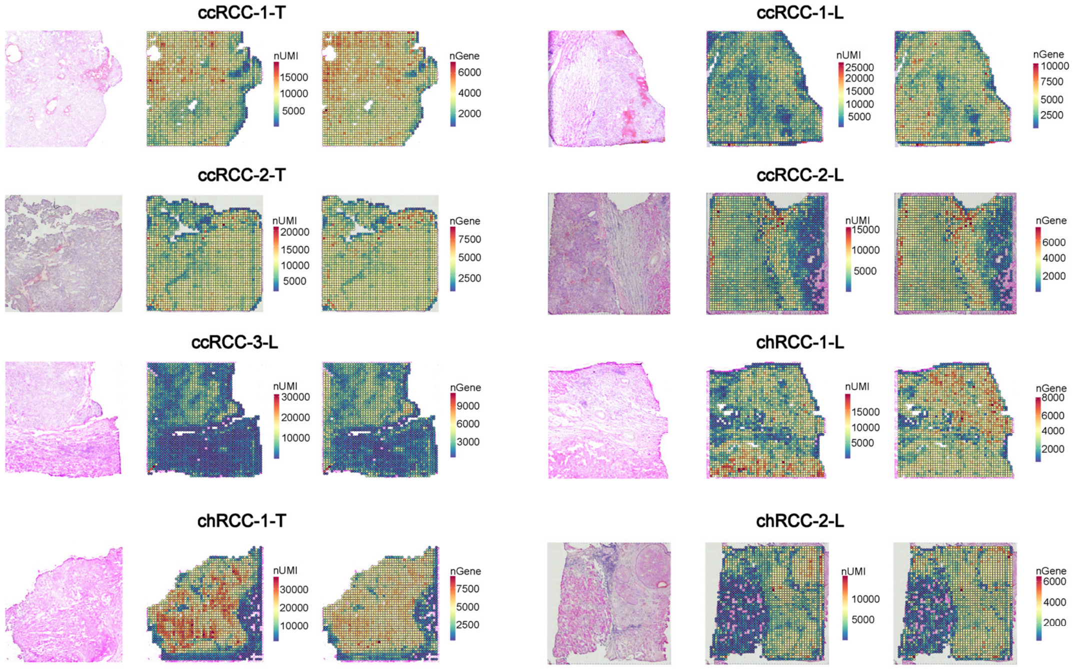

**Supplementary Figure S17. Evaluation of RCC data quality.** Columns 1 and 4, tissue sections; columns 2 and 5, spatial feature plots showing the number of UMIs (nUMI); columns 3 and 6, spatial feature plots showing the number of genes (nGene).

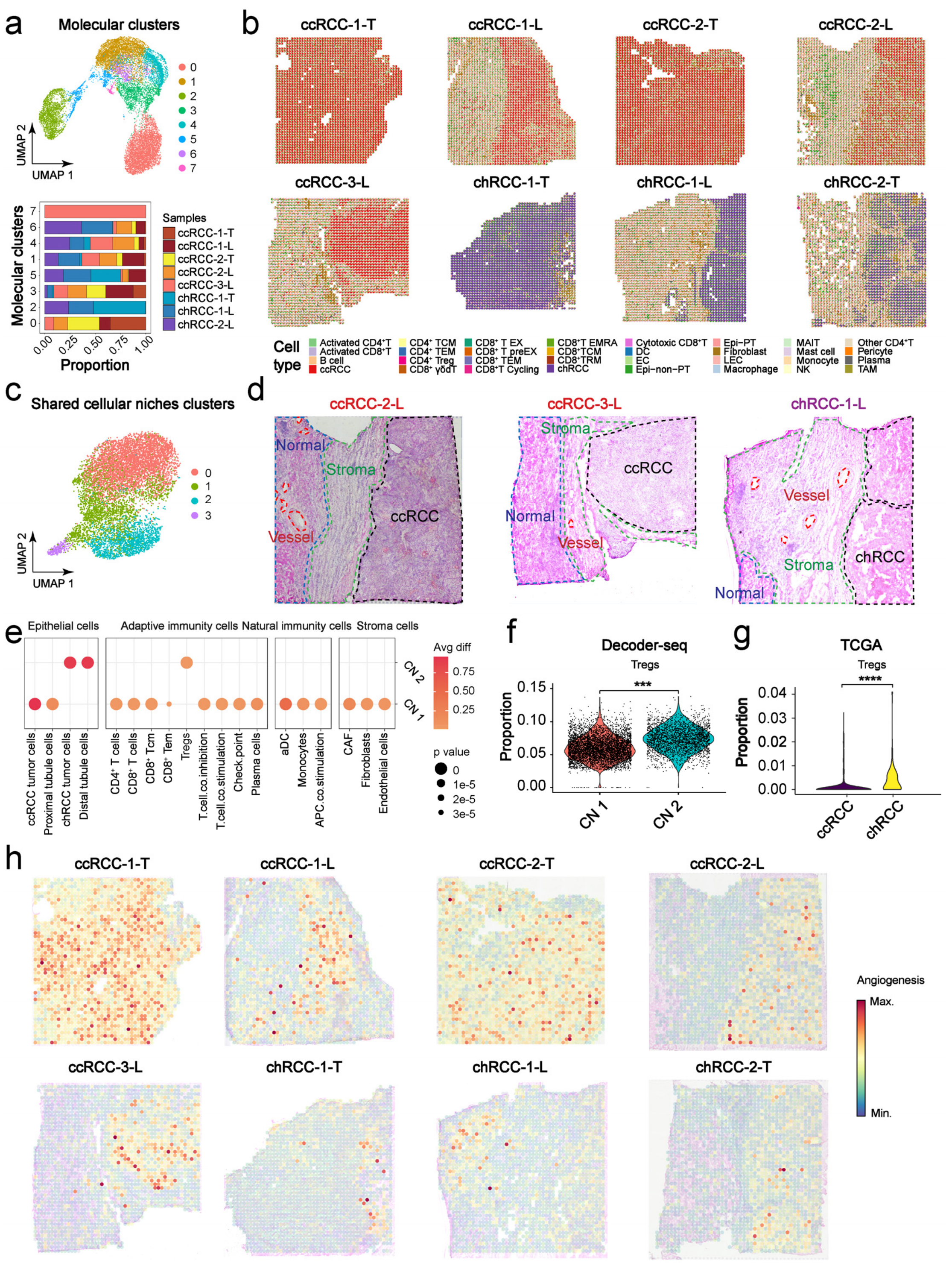

**Supplementary Figure S18. Resolving the spatial heterogeneity in RCCs with Decoder-seq. Related to Figure 5. (a)** Unsupervised clustering of Decoder-seq RCC data (upper panel); the proportion of each cluster in each sample (lower panel). **(b)** Deconvolution of cell type composition. **(c)** Cellular niche (CN) clustering in the shared cluster. Epi-non-PT: Epithelial-non-proximal tubule cell; Epi-PT: Epithelial-proximal tubule cell; LEC: Lymphatic endothelial cell; EC: endothelial cell; TAM:

tumor-associated macrophage; $CD8^+$ T EMRA: $CD8^+$ terminally differentiated effector memory (T EMRA) cell; $CD8^+$/$CD4^+$ TCM: $CD8^+$/$CD4^+$ T central memory cell; $CD8^+$/$CD4^+$ TEM: $CD8^+$/$CD4^+$ T effector memory cell; NK: natural killer cell; DC: dendritic cell; $CD8^+$ TRM: $CD8^+$ tissue-resident memory cell; $CD8^+$ T EX: Exhausted CD8+ T cell; $CD8^+$ T preEX: Precursor exhausted $CD8^+$ T cell. MAIT: Mucosal-associated invariant T cell. **(d)** H&E images of ccRCC-2-L, ccRCC-3-L, and chRCC-1-L. Pathological annotation of morphological regions into distinct categories including normal renal (dark blue), stromal (green), blood vessel (red), and renal cell carcinoma (black). **(e)** Bubble diagram showing differential analysis of CN 1 and CN 2 based on cell scoring. **(f)** Comparison of regulatory T cell (Treg) abundance in CN 1 and CN 2. **(g)** Comparison of Treg abundance in ccRCC and chRCC subtype samples based on data from The Cancer Genome Atlas (TCGA). **(h)** Spatial projection of the angiogenic pathway in each tissue section.

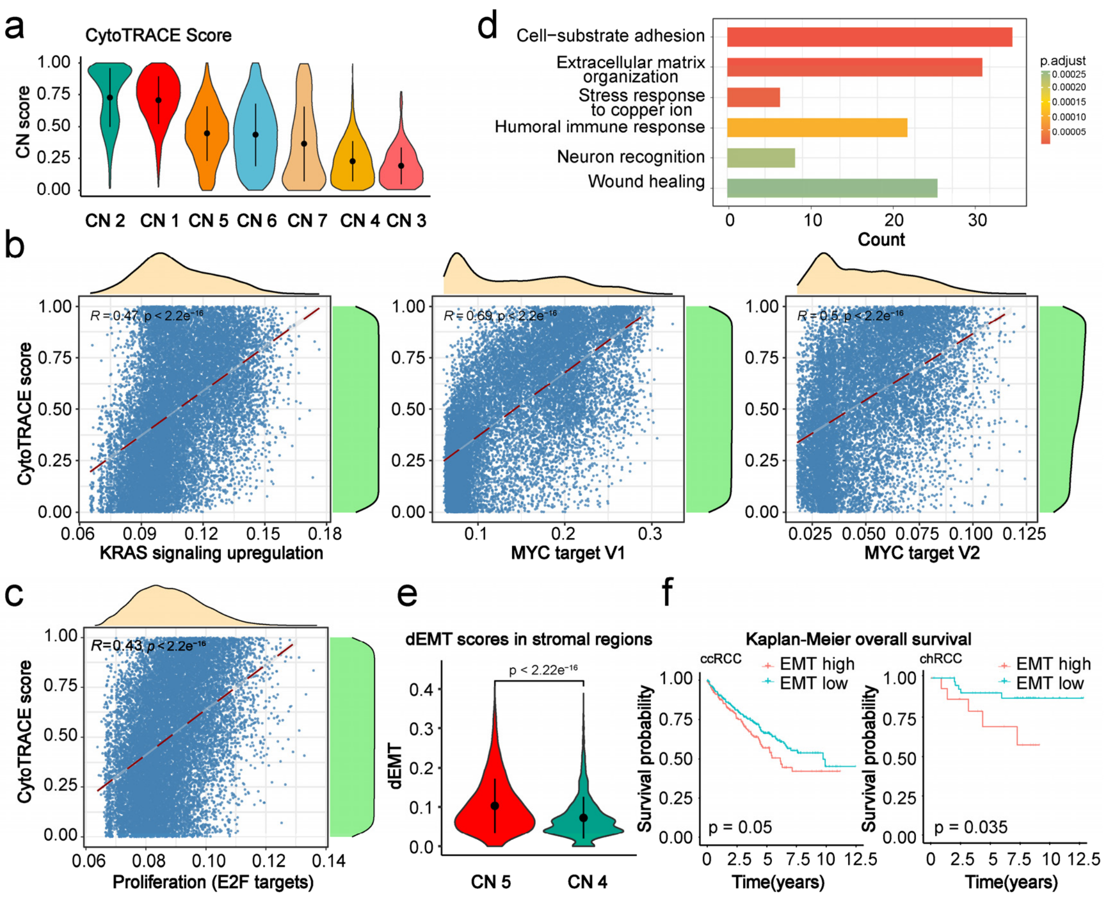

**Supplementary Figure S19. Deciphering the spatial evolutionary trajectory of tumor invasion in RCCs. Related to Figure 6. (a)** CytroTRACE scores in CNs. **(b)** Spearman's correlation analysis between CytroTRACE scores and KRAS signaling upregulation, MYC targets V1, and V2 from the HALLMARK gene sets for each spot. **(c)** Spearman's correlation analysis between CytroTRACE scores and cell proliferation related-E2F targets for each spot. **(d)** Gene Ontology term analysis of spatially upregulated signatures (SUSs). **(e)** Comparison of dynamic Epithelial-mesenchymal transition (EMT) (dEMT) in stromal regions between CN 4 and CN 5. **(f)** Survival curves and primary T stages of cases with high and low HALLMARKS EMT scores based on data from TCGA. Statistical significance was assessed with a log-rank test.

**Supplementary Table S1. Comparison of spatial DNA barcodes (oligo-dT numbers) per $\mu m^2$ for substrates used in ST[5] and Pixel-seq[6], 2D-homemade slide, 2D commercial slide, and our 3D dendrimeric slide.**

| | Array substrate | oligo-dT numbers per $\mu m^2$ |
|---|---|---|
| ST | 3D linear NHS modification | $2.55\times10^4$ |
| Pixel-seq | Cross-linked polyacrylamide coating | $2.03\times10^4$ |
| 2D-homemade | 2D linear amino modification | $1.16\times10^4$ |
| 2D commercial | 2D linear amino modification | $1.46\times10^4$ |
| Decoder-seq | 3D dendrimer amino modification | $2.04\times10^5$ |

**Supplementary Table S2. Cost object breakdown of 3D dendrimeric slide.**

| 3D dendrimeric slide | SOURCE | IDENTIFIER | Price (CNY) | Cost (CNY)/Per slide |
|---|---|---|---|---|
| Glass slide | SAIL BRAND | 7101 | 8.00 | 0.16 |
| GOPTS | Sigma | 440167 | 610.00 | 2.20 |
| Ethanol | Shanghai Lingfeng | 64-17-5 | 18.50 | 0.12 |
| PAMAM | Sigma | 412449 | 7,717 | 44.00 |
| Methanol | Shanghai Lingfeng | 67-56-1 | 24.00 | 0.12 |
| **Total（per 3D dendrimeric slide）** | | | | **￥46.6/$6.7** |

**Supplementary Table S3. Cost object breakdown of DNA coordinate array.**

| DNA coordinate array | SOURCE | IDENTIFIER | Price (CNY) | Cost (CNY)/Per slide |
|---|---|---|---|---|
| PDMS | CChip | LHC001 | 1200 | 10 |
| DNA barcode X | Songon | NA | 12000 | 9 |
| DNA barcode Y | Songon | NA | 14625 | 12.83 |
| Linker | Songon | NA | 106 | 4.05 |
| T4 DNA ligase | Vazyme | C301-01 | 96 | 6 |
| Succinic anhydride | Aladdin | S104823 | 24 | 0.12 |
| Triethylamine | Macklin | T818772 | 118 | 0.12 |
| DMF | Sinopharm | H1E0140 | 44 | 0.88 |
| DSS | Thermo Fisher Scientific | 21655 | 2654 | 0.88 |
| **Total（per DNA coordinate array）** | | | | **￥43.88/$6.1** |

**Supplementary Table S4. Mean and standard deviation of gene measurements in Figure. 3c.**

| Genes | Decoder-seq | ISS |
|---|---|---|
| *Fabp7* | 16426.33 ± 4484.81 | 41894.33 ± 4902.42 |
| *Penk* | 13152.67 ± 2654.91 | 26892.00 ± 4104.96 |
| *Slc17a7* | 21684.67 ± 4789.16 | 18423.00 ± 2016.27 |

**Supplementary Table S5. Mean and standard deviation of lowly-expressed *Olfr*-gene measurements in Figure. 3h.**

| Method | Type | Number |
|---|---|---|
| Decoder-seq (50 μm) | 757.67 ± 126.18 | 2468.67 ± 698.32 |
| 10× Visium (55 μm) | 132 ± 22.12 | 461 ± 39.69 |

**Supplementary Table S6. List of all *Olfr*-genes identified in 10× Visium data and Decoder-seq data.**

**Supplementary Table S7. Dendritically enriched gene-sets.**

Sheet 1: Dendritic-enriched genes identified by Decoder-seq along with Fold-Change enrichment, and FDR corrected q-value ($p < 0.05$, logFC > 0.8).
Sheet 2: Soma-enriched genes identified by Decoder-seq along with Fold-Change enrichment, and FDR corrected q-value ($p < 0.05$, logFC < -0.8).
Sheet 3: List of genes that overlap with Aingsley et al., 2014[1], Tushev et al., 2018[2], Nakayama et al., 2017[3], and Slide-seqV2[4].

**Supplementary Table S8. RCC data quality evaluation.**

| Sample | RIN | Average genes per Spot | Average UMI Counts per Spot | Sequencing saturation |
|---|---|---|---|---|
| ccRCC-1-T | 6.36 | 3930 | 8956 | 83% |
| ccRCC-1-L | 7.35 | 3763 | 6958 | 84% |
| ccRCC-2-T | 5.27 | 3955 | 8034 | 83% |
| ccRCC-2-L | 8.3 | 3009 | 4870 | 91% |
| ccRCC-3-L | 5.28 | 2257 | 3975 | 89% |
| chRCC-1-T | 3.32 | 4577 | 17935 | 61% |
| chRCC-1-L | NA | 3492 | 6721 | 79% |
| chRCC-2-L | 5.25 | 1996 | 3664 | 93% |

**Supplementary Table S9. List of cell signatures for cell scoring.**

**Supplementary Table S10. List of spatially upregulated genes.**

**Supplementary Table S11. Go enrichment analysis of spatially upregulated genes.**

**Supplementary Table S12. List of dynamic EMT genes.**

**Supplementary Table S13. DNA sequences.**

| Name | Sequence |
|---|---|
| DNA barcode X | 5’ $NH_2$-C6-CTACACGACGCTCTTCCGATCT [barcode X] AGGCCAGAGCATTCG 3’ |
| DNA barcode Y | 5’ Phos- ATCCACGTGCTTGAG [barcode Y] NNNNNNNNNNNNTTTTTTTTTTTTTTTTTTTTTTTTTTTTTTVN |
| Ligation linker | CTCAAGCACGTGGATCGAATGCTCTGGCCT |
| TSO | AAGCAGTGGTATCAACGCAGAGTACATrGrGrG |
| Second strand primer | AAGCAGTGGTATCAACGCAGAGTGANNNGGNNNB |
| Forward primer | CTACACGACGCTCTTCCGATCT |
| Reverse primer | AAGCAGTGGTATCAACGCAGAG |
| i5 primer | AATGATACGGCGACCACCGAGATCTACAC[i5] ACACTCTTTCCCTACACGACGCTC |

### References

1. Ainsley, J.A., Drane, L., Jacobs, J., Kittelberger, K.A. & Reijmers, L.G. Functionally diverse dendritic mRNAs rapidly associate with ribosomes following a novel experience. *Nat. Commun.* **5**, 4510 (2014).
2. Tushev, G. et al. Alternative 3' UTRs Modify the Localization, Regulatory Potential, Stability, and Plasticity of mRNAs in Neuronal Compartments. *Neuron* **98**, 495-511 e496 (2018).
3. Nakayama, K. et al. RNG105/caprin1, an RNA granule protein for dendritic mRNA localization, is essential for long-term memory formation. *eLife* **6**, e29677 (2017).
4. Stickels, R.R. et al. Highly sensitive spatial transcriptomics at near-cellular resolution with Slide-seqV2. *Nat. Biotechnol.* **39**, 313-319 (2021).
5. Ståhl, P.L. et al. Visualization and analysis of gene expression in tissue sections by spatial transcriptomics. *Science* **353**, 78-82 (2016).
6. Fu, X. et al. Polony gels enable amplifiable DNA stamping and spatial transcriptomics of chronic pain. *Cell* **185**, 4621-4633 e4617 (2022).
